# Structural variants in the barley gene pool: precision and sensitivity to detect them using short-read sequencing and their association with gene expression and phenotypic variation

**DOI:** 10.1101/2022.04.25.489331

**Authors:** Marius Weisweiler, Christopher Arlt, Po-Ya Wu, Delphine Van Inghelandt, Thomas Hartwig, Benjamin Stich

## Abstract

In human genetics, several studies have shown that phenotypic variation is more likely to be caused by structural variants (SV) than by single nucleotide variants (SNV). However, accurate while cost-efficient discovery of SV in complex genomes remains challenging. The objectives of our study were to (i) facilitate SV discovery studies by benchmarking SV callers and their combinations with respect to their sensitivity and precision to detect SV in the barley genome, (ii) characterize the occurrence and distribution of SV clusters in the genomes of 23 barley inbreds that are the parents of a unique resource for mapping quantitative traits, the double round robin population, (iii) quantify the association of SV clusters with transcript abundance, and (iv) evaluate the use of SV clusters for the prediction of phenotypic traits. In our computer simulations based on a sequencing coverage of 25x, a sensitivity *>*70% and precision *>*95% was observed for all combinations of SV types and SV length categories if the best combination of SV callers was used. We observed a significant (P *<* 0.05) association of gene-associated SV clusters with global gene-specific gene expression. Furthermore, about 9% of all SV clusters that were within 5kb of a gene were significantly (P *<* 0.05) associated with the gene expression of the corresponding gene. The prediction ability of SV clusters was higher compared to that of single nucleotide polymorphisms from an array across the seven studied phenotypic traits. These findings suggest the usefulness of exploiting SV information when fine mapping and cloning the causal genes underlying quantitative traits as well as the high potential of using SV clusters for the prediction of phenotypes in diverse germplasm sets.

## INTRODUCTION

Researchers began to study genomic rearrangements and structural variants (SV) about 60 years ago. These studies investigated somatic chromosomes, biopsies, and cell cultures from lymphomas to understand the role of abnormal chromosome numbers as well as SV for the development of cancer (Jacobs and Strong, 1959; Nowell and Hungerford, 1960; Manolov and Manolov, 1972; Craig-Holmes et al., 1973; Mitelman et al., 1979).

The development of sequencing by synthesis pioneered by Frederick Sanger (Sanger et al., 1977) enabled in the following years the first sequenced genomes of prokaryotes (e.g. *Escherichia coli* ) and eukaryotes (e.g. yeast) (Goffeau et al., 1996; Blattner et al., 1997). Next milestones of sequencing by synthesis were the sequenced genomes of *Arabidopsis thaliana* as first plant species (The Arabidopsis Genome Initiative, 2000) and of human (Craig Venter et al., 2001). Due to the development of next-generation sequencing (NGS) platforms such as 454 and Illumina, studies aiming for genome-wide variant detection in 100s or 1000s of samples as in the 1000 genome project (Altshuler et al., 2012) became possible.

Three different approaches have been proposed to detect SV based on NGS data: assembling, long-read sequencing, and short-read sequencing (Mahmoud et al., 2019). For crop and especially for cereal species, the assembly approach is a tough challenge because of the large genome size and the high proportion of repetitive elements in the genomes (Neale et al., 2014; Mascher et al., 2017). Long-read mapping requires Pacific Biosciences or Nanopore sequencing data which results in high costs if many accessions should be sequenced and, thus, is not affordable for many research groups.

In contrast, short-read sequencing is well-established for SV detection in the human genome (Chaisson et al., 2019; Ebert et al., 2021). Various software tools have been developed to detect SV from short-read sequencing data and were benchmarked based on human genomes (Cameron et al., 2019; Kosugi et al., 2019).

More recently there is also an increased interest in using such approaches for SV detection in plant genomes (Fuentes et al., 2019; Zhou et al., 2019; Guan et al., 2021). Fuentes et al. (2019) evaluated several SV callers to detect SV in the rice genome. However, no study evaluated the performance of SV callers for transposon-rich complex cereal genomes.

Several studies have examined the distribution and frequency of SV in the genomes of rice and maize (Wang et al., 2018; Yang et al., 2019; Kou et al., 2020). Despite the importance of cereals for human nutrition, only Jayakodi et al. (2020) performed a genome-wide study on SV in barley, with a focus on large SV in 20 barley accessions. In humans, SV have been described to have an up to ∼50fold stronger influence on gene expression than single nucleotide variants (SNV) (Chiang et al., 2017). SV also have been associated with changes in transcript abundance in plants such as in cucumber (Zhang et al., 2015), maize (Yang et al., 2019), tomato (Alonge et al., 2020), and soybean (Liu et al., 2020a). However, the role and frequency of SV in gene regulatory mechanisms in small grain cereals is widely unexplored.

In humans, several studies have shown that phenotypic variation is more likely to be caused by SV than by SNV (Alkan et al., 2011; Baker, 2012; Sudmant et al., 2015; Schüle et al., 2017; McColgan and Tabrizi, 2018). In plants, individual SV have been associated with traits such as Aluminium tolerance in maize (Maron et al., 2013), disease resistance and domestication in rice (Xu et al., 2012), or plant height (Li et al., 2012) and heading date (Nishida et al., 2013) in wheat. In barley, individual SV have been associated with traits such as Boron toxicity tolerance (Sutton et al., 2007) and disease resistance (Muñoz-Amatriaín et al., 2013). In grapevine and rice, it has been shown that SV have a low variant frequency due to purifying selection (Zhou et al., 2019; Kou et al., 2020). However, few studies have examined the ability to predict quantitatively inherited phenotypic traits using SV in comparison to SNV.

The objectives of our study were to (i) facilitate SV discovery studies by benchmarking SV callers and their combinations with respect to their sensitivity and precision to detect SV in the barley genome, (ii) characterize the occurrence and distribution of SV clusters in the genomes of 23 barley inbreds that are the parents of a unique resource for mapping quantitative traits, the double round robin population (Casale et al., 2021), (iii) quantify the association of SV clusters with transcript abundance, and (iv) evaluate the use of SV clusters for the prediction of phenotypic traits.

## RESULTS

### Precision and sensitivity of SV callers

Six tools (Table 1) which call SV based on short-read sequencing data were evaluated with respect to their precision and sensitivity to detect five different SV types (deletions, insertions, duplications, inversions, and translocations) in five SV length categories (A: 50 - 300bp; B: 0.3 - 5kb; C: 5 - 50kb; D: 50 - 250kb; E: 0.25 - 1Mb) using computer simulations. The precision of Delly, Manta, GRIDSS, and Pindel to detect deletions of all five SV length categories based on 25x sequencing coverage ranged from 97.8 - 100.0%, whereas the precision of Lumpy and NGSEP was lower with values between 75.0 and 89.8% (Supplementary Table 2). The sensitivity of NGSEP was with 78.6 - 87.5% the highest but that of Manta was with 79.7 - 81.1% only slightly lower. We evaluated various combinations of SV callers and observed for the combination of Manta | GRIDSS | Pindel | Delly | (Lumpy & NGSEP) an increase of the sensitivity to detect deletions compared to the single SV callers up to a final of 89.0% without decreasing the precision considerably (99.1%).

**Table 1:**
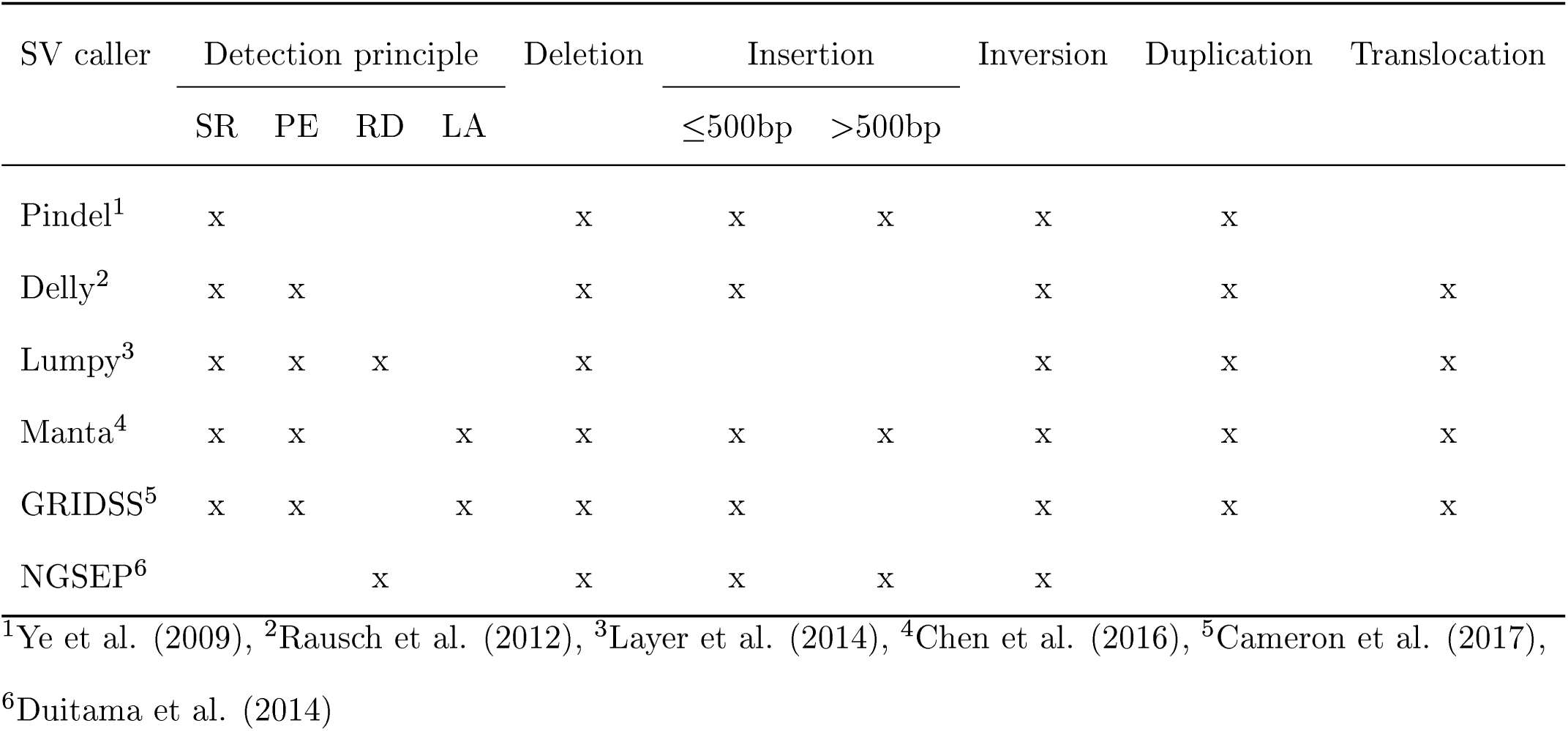
Properties of structural variant (SV) callers for short-read sequencing that were compared in our study, where split reads (SR), paired-end reads (PE), read depth (RD), and local alignments (LA) are the underlying detection principles.

Manta was the only SV caller which allowed the detection of insertions of all SV length categories with precision values as high as 99.8 to 100.0%. The combination of Manta | GRIDSS | Delly for the SV length category A has shown a high sensitivity (88.4%) and precision (99.8%). This combination was therefore used for the detection of insertions of SV length category A in further analyses.

The sensitivity of the SV callers Delly, Manta, Lumpy, and GRIDSS to detect duplications of the SV length category A was with values from 28.2 to 39.4% very low. In contrast, Pindel could detect these duplications with a sensitivity of 75.7%. For the other SV length categories, the combination of Manta | GRIDSS | Pindel could increase the sensitivity to detect duplications by 2 to 7% compared to using a single SV caller while the precision ranged between 97.6 and 99.3%.

The performance of Lumpy and NGSEP to detect inversions reached precision values of 81.5 - 98.5% and sensitivity values of 66.1 - 80.0% that were on the same low level as for deletions. Delly performed well for detecting inversions in SV length categories B to D, but for E and especially for A, the performance was lower compared to that of the other SV callers. Overall, Pindel was the only SV caller with a combination of both, high precision and sensitivity to detect inversions. These precision and sensitivity values could be further improved across all SV length categories by combining the calls of Pindel with that of Manta | GRIDSS (Supplementary Table 2).

The combination of GRIDSS | Pindel | GATK increased the sensitivity to detect small insertions and deletions (2 - 49bp, INDELs) by 3% compared to using the single callers (Supplementary Table S1). With 6%, an even higher difference for the sensitivity to detect translocations was observed between the combination of Manta | GRIDSS | (Delly & Lumpy) and single callers.

In a next step, different sequencing coverages from 1.5x to 65x were simulated and the performance of the best combination of SV callers for each of the SV types was compared to their performance with 25x sequencing coverage (Supplementary Fig. S1). For deletions, the F1-score, which is harmonic mean of the precision and sensitivity, for 65x sequencing coverage was ∼2% higher than for 25x sequencing coverage. Only marginal differences were observed between the F1-score of 65x or 25x sequencing coverage for calling duplications and inversions. Interestingly, the F1-score for calling translocations and insertions was with 2% and 9%, respectively, higher in the scenario with 25x than with 65x sequencing coverage. For 12.5x sequencing coverage, the F1- score was still on an high level with values *>* 80% for each SV type (Supplementary Fig. S2). With a further reduced sequencing coverage, the F1-score also decreased. Finally, the performance of our pipeline to detect SV was evaluated based on 14x and 25x linked-read sequencing data. For all SV types and SV length categories, with the exception of deletions and duplications in SV length category D and A, respectively, the F1-score was 2 to 7% higher based on Illumina sequencing data than based on linked-read sequencing data.

### SV clusters across the 23 parental inbreds of the double round robin population

Across the 23 barley inbreds, that are the parents of a new resource for mapping natural phenotypic variation, the double round robin population, we detected 458,671 SV clusters using the best combination of SV callers (Table 3). These comprised 183,489 deletions, 70,197 insertions, 93,079 duplications, 6,583 inversions, and 105,323 translocations. Additionally, 12,734,736 INDELs were detected across the seven chromosomes. The proportion of SV clusters which were annotated as transposable elements varied from 1.4% for inversions to 51.5% for translocations.

**Table 2:** Sensitivity/precision of structural variant (SV) callers and combinations of them (for details see Material & Methods) to detect deletions, insertions, duplications, and inversions of the SV length categories A (50 - 300bp), B (0.3 - 5kb), C (5 - 50kb), D (50 - 250kb), and E (0.25 - 1Mb).

| SV caller | SV length category |  |  |  |  |
| --- | --- | --- | --- | --- | --- |
|  | A | B | C | D | E |
| Deletions |  |  |  |  |  |
| Delly | 58.1/97.8 | 76.2/99.4 | 72.5/99.3 | 72.4/100.0 | 75.0/100.0 |
| Manta | 79.7/100.0 | 81.1/99.8 | 79.9/99.6 | 79.7/99.4 | 81.0/100.0 |
| Lumpy | 60.0/78.1 | 70.5/86.5 | 66.8/85.6 | 62.5/79.0 | 64.3/80.6 |
| GRIDSS | 79.0/99.5 | 80.7/99.9 | 77.8/99.9 | 78.1/100.0 | 77.4/100.0 |
| Pindel | 87.4/99.9 | 68.4/99.7 | 83.6/99.4 | 80.2/100.0 | 67.9/100.0 |
| NGSEP | 84.1/87.3 | 83.1/83.4 | 83.5/82.2 | 87.5/89.8 | 78.6/75.0 |
| Combination | 89.0/99.1 | 86.9/99.4 | 86.7/99.2 | 86.5/99.4 | 86.9/100.0 |
| Insertions |  |  |  |  |  |
| Delly | 3.4/100.0 |  |  |  |  |
| Manta | 88.4/99.8 | 74.1/100.0 | 72.1/100.0 | 72.5/100.0 | 75.0/100.0 |
| GRIDSS | 45.5/100.0 |  |  |  |  |
| Pindel | 6.6/93.0 |  |  |  |  |
| NGSEP | 64.1/59.2 | 26.8/29.6 | 35.5/40.5 | 30.5/32.1 | 26.0/26.5 |
| Combination | 88.4/99.8 | 74.1/100.0 | 72.1/100.0 | 72.5/100.0 | 75.0/100.0 |
| Duplications |  |  |  |  |  |
| Delly | 28.2/99.0 | 75.1/96.8 | 74.7/95.4 | 75.3/97.2 | 71.7/91.7 |
| Manta | 39.0/99.5 | 80.5/99.8 | 82.7/99.8 | 83.9/98.7 | 82.6/97.4 |
| Lumpy | 31.5/98.4 | 67.9/84.8 | 67.7/82.6 | 68.3/81.9 | 65.2/80.0 |
| GRIDSS | 39.4/99.8 | 80.0/100.0 | 80.0/100.0 | 83.3/100.0 | 79.4/100.0 |
| Pindel | 75.7/98.1 | 57.8/99.0 | 88.1/99.8 | 83.9/99.4 | 73.9/100.0 |
| Combination | 75.8/98.1 | 87.3/99.1 | 90.8/99.3 | 89.8/98.2 | 89.1/97.6 |
| Inversions |  |  |  |  |  |
| Delly | 49.7/70.4 | 84.6/99.2 | 85.5/99.4 | 82.6/99.4 | 78.2/98.6 |
| Manta | 77.0/99.0 | 87.0/99.9 | 87.3/99.9 | 90.0/100.0 | 82.8/100.0 |
| Lumpy | 66.1/88.5 | 76.8/96.2 | 75.3/97.4 | 77.4/94.8 | 74.7/98.5 |
| GRIDSS | 76.9/99.1 | 86.9/99.8 | 85.2/99.9 | 87.9/100.0 | 82.8/100.0 |
| Pindel | 83.5/99.2 | 90.7/99.9 | 90.2/99.9 | 89.0/100.0 | 77.0/100.0 |
| NGSEP | 0.0/0.0 | 75.7/87.9 | 75.3/81.5 | 80.0/85.4 | 77.0/88.2 |
| Combination | 88.4/98.1 | 91.5/99.8 | 90.9/99.8 | 93.2/100.0 | 85.1/100.0 |

**Table 3:** Summary of detected structural variants (SV) and small insertions and deletions (2 - 49bp, INDELs) across 23 diverse barley inbreds, where MAF was the minor allele frequency, and TE were SV clusters which were annotated as transposable elements in the Morex reference sequence v3.

| SV type | Number of SV calls | Number of SV clusters |  |  |
| --- | --- | --- | --- | --- |
|  |  |  | MAF > 0.05 | TE |
| Deletions | 714,867 | 183,489 | 78,823 | 16,846 |
| Insertions | 241,522 | 70,197 | 29,672 | 279 (17,718)<sup>1</sup> |
| Duplications | 195,710 | 93,079 | 58,793 | 6,608 |
| Inversions | 14,961 | 6,583 | 4,116 | 92 |
| Translocations | 251,956 | 105,323 | 61,572 | 0 (54,258)<sup>1</sup> |
| INDELs | 59,934,113 | 12,734,736 | 4,492,832 | 32 |

<sup>1</sup>Because of missing endpoint information no reciprocal overlap criterion applied

We performed a PCR based validation for detected deletions and insertions (Supplementary Table S2, Supplementary Fig. S3). Six out of six deletions and five out of five insertions up to 0.3kb could be validated (Supplementary Fig. S4). Additionally, we could validate eight out of eleven deletions between 0.3kb and 460kb (Supplementary Fig. S5), where for the three not validated deletions, the expected fragments were not observed in the non-reference parental inbred.

The number of SV clusters present per inbred ranged from less than 40,000 to more than 80,000 (Fig. 1A). We observed no significant (P *>* 0.05) correlation between the sequencing coverage, calculated based on raw, trimmed, and mapped reads, of each inbred as well as the number of detected SV clusters in the corresponding inbred. A two-sided t-test resulted in no significant (P *>* 0.05) association between the number of SV clusters of an inbred and the spike morphology as well as the landrace vs. variety status of the inbreds. In contrast, principal component analyses based on presence/absence matrices of the SV clusters revealed a clustering of inbreds by spike morphology, geographical origin, and landrace vs. variety status (Supplementary Fig. S6).

**Fig. 1:**
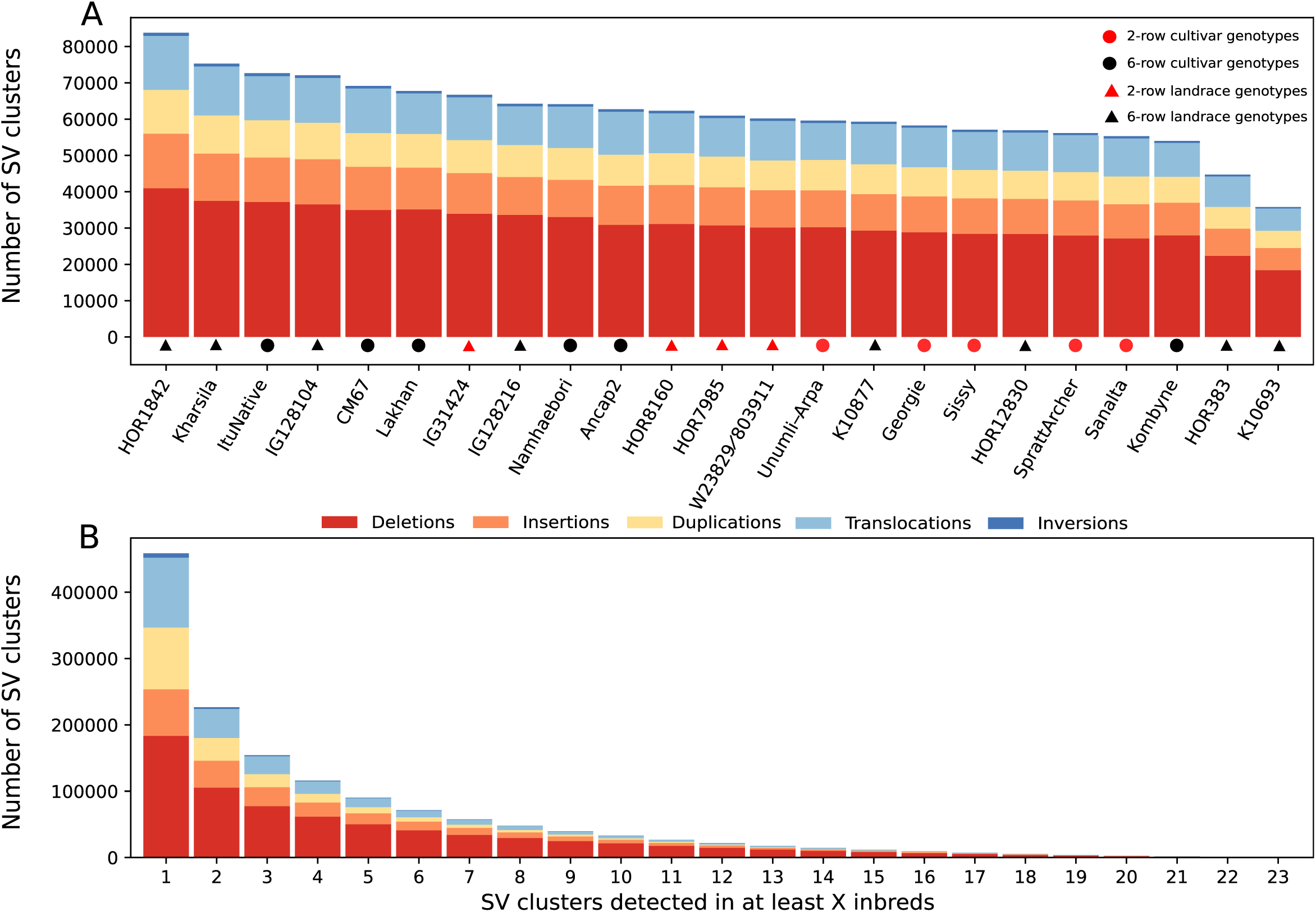
Stacked bar graph of the number of different types of structural variant (SV) clusters detected in the 23 inbreds (A) and SV clusters which were detected in at least the given number of the inbreds (B).

Out of the 458,671 SV clusters, 50.6% (232,071) appeared in only one of the 23 inbreds, whereas 19.7% (90,256) were detected in at least five inbreds (Fig. 1B, Supplementary Fig. S7). Additional analyses revealed a significant although weak negative correlation (r = −0.06681, P = 2.07x10^−314^) between the length of a SV cluster and its minor allele frequency (MAF). The average MAF of SV clusters with a length of 250kb to 1Mb and of 50 - 250kb was 0.08, respectively, while that of SV clusters with a length of 50bp - 50kb was 0.13 (Supplementary Fig. S8). SV clusters annotated as transposable elements had a shorter average length of 5,853bp and a higher MAF of 0.16 compared to SV clusters that were not annotated as transposable elements (10,605bp, 0.12). Deletions and insertions of the SV length category A were the most common detected SV clusters with a fraction of 41.7 and 48.4%, respectively (Supplementary Table S3). In contrast, for duplications, the largest fraction were that for SV clusters of the SV length category C (55.9%). The average MAF of the individual SV types was the highest for insertions with 0.17, followed by deletions, inversions, translocations, and duplications with values of 0.14, 0.11, 0.10, and 0.10, respectively.

### Characterization of the SV clusters

After examining the length of the detected SV clusters and their presence in the 23 barley inbreds, we investigated the distribution of the SV clusters across the barley genome. We observed a significant correlation (r = 0.5653, P *<* 0.01) of nucleotide diversity (*π*) of SV clusters and SNV, measured in 100kb windows along the seven chromosomes (Supplementary Fig. S9). The SV clusters were predominantly present distal of pericentromeric regions. In contrast to SNV, the frequency of all SV types, and especially that of duplications, increased in centromeric regions (Fig. 2). For all centromeres, a significantly (P *<* 0.01) higher number of SV clusters was observed compared to what is expected based on a poisson distribution and, thus, were designated as SV hotspots. The proportion of SV clusters in pericentromeric regions was with 14.5% considerably lower compared to what is expected based on the physical length of these regions (25.7%). Only 4.5% of all detected SV hotspots were observed in pericentromeric regions. Compared to the five SV types, the genome-wide distribution of INDELs was more equal. Their occurrences peaked not only within, but also distal to pericentromeric and centromeric regions.

**Fig. 2:**
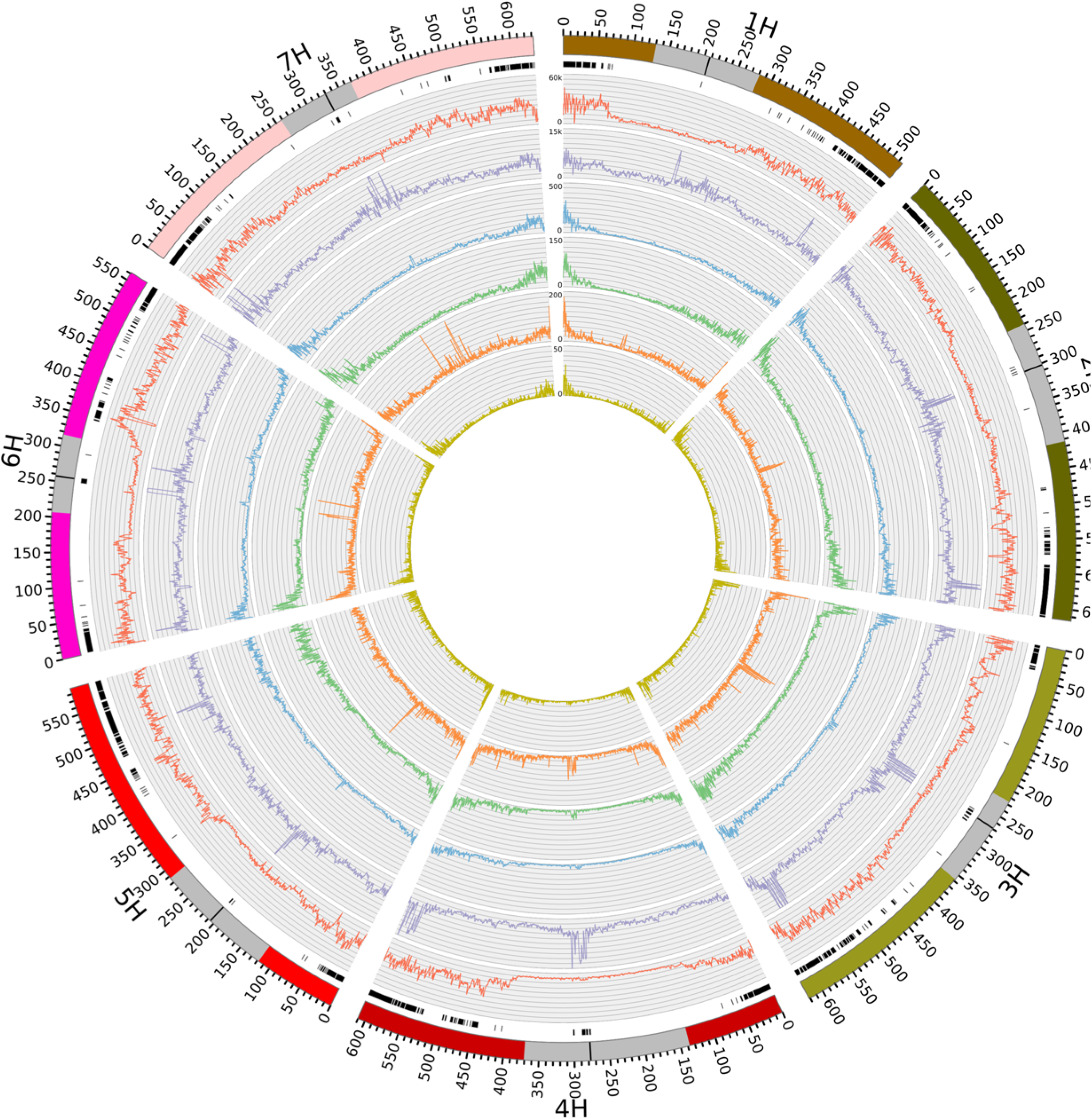
Distribution of genomic variants among 23 barley inbreds across the seven chromosomes. The outermost circle denotes the chromosome number, the physical position, and as gray bar the peri-centromeric regions (Casale et al. 2021) plus the centromeres (black) according to the Morex reference sequence v3. The next inner circles report the SV cluster hotspots (black bars), frequencies of single nucleotide variants (red), small insertions and deletions (2 - 49bp, INDELs, purple), deletions (blue), insertions (green), duplications (orange), and inversions (yellow) which were detected among the 23 inbreds.

We also examined if SV clusters provide additional genetic information compared to that of closely linked SNV. To do so, we determined the extent of linkage disequilibrium (LD) between each SV cluster and SNV located within 1kb and compared this with the extent of LD between the closest SNV to the SV cluster and the SNV within 1kb. Across the different SV types, 33.7 - 74.3% have at least one SNV within 1kb that showed an r^2^ ≥ 0.6 (Supplementary Table S4). In contrast, 89.2 - 89.9% of SNV that are closest to the SV cluster showed an r^2^ ≥ 0.6 to another SNV within 1kb. In the next step, we examined the presence of SV clusters relative to the position of genes. The highest proportion of SV clusters (∼ 60%) was located in intergenic regions of the genome (Fig. 3). The second largest fraction (∼ 30%) of SV clusters was present in the 5kb up- or downstream regions of genes, which is considerably higher compared to that of INDELs (∼ 17%) and SNV (∼ 16%). Within the group of SV clusters that were 5kb up- or downstream to genes, a particularly high fraction were inversions. On average across all SV types, about 10% of SV clusters were located in introns and exons, with inversions being the exception again, showing a considerably higher rate.

**Fig. 3:**
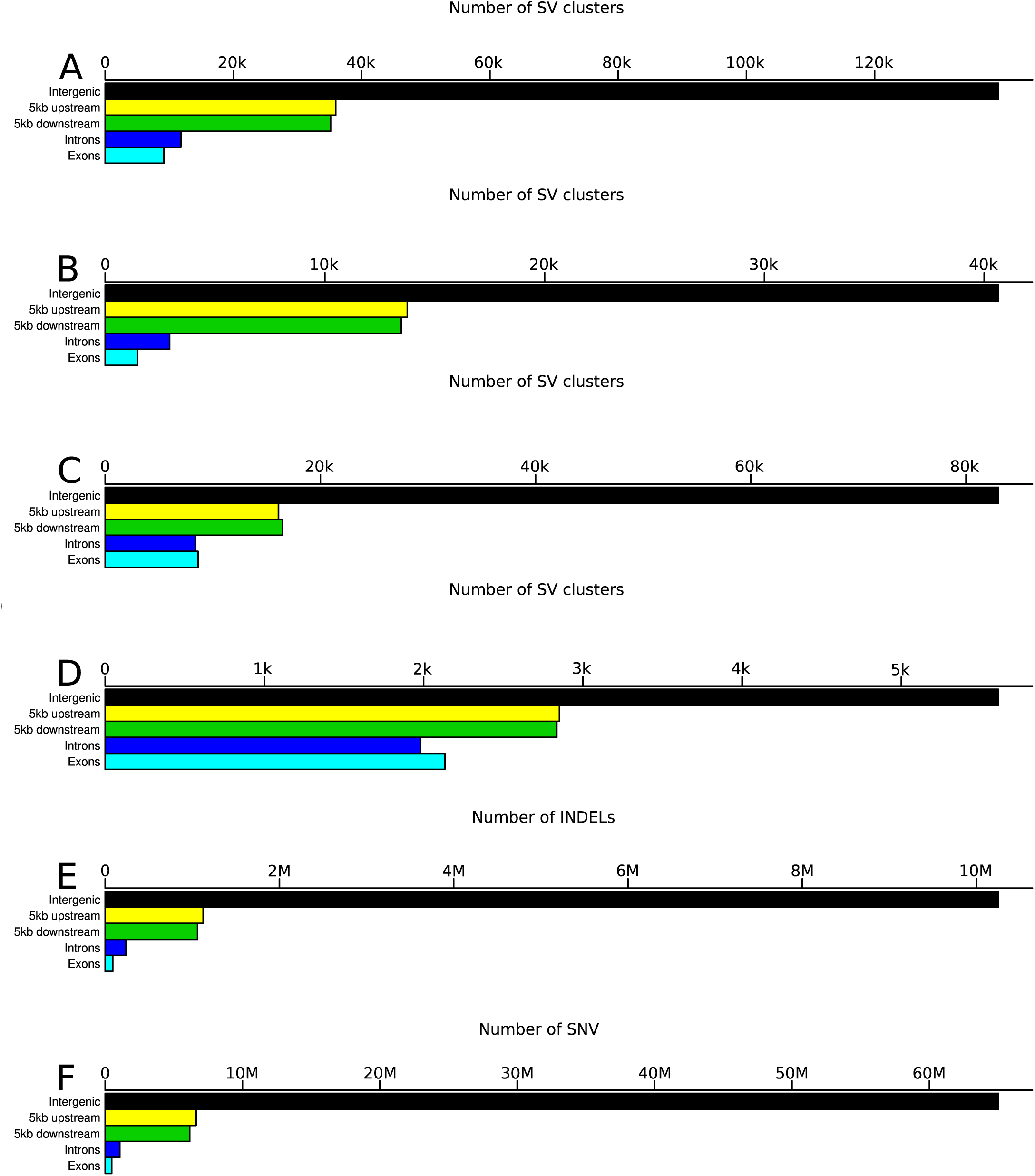
The occurrence of deletions (A), insertions (B), duplications (C), inversions (D), small insertions and deletions (2 - 49bp, INDELs, E), and single nucleotide variants (SNV) (F) in five genomic regions.

The enrichment of SV clusters proximal to genes lead us to assess their physical distance relative to the transcription start site (TSS) of the closest genes and compare this to SNV. The number of SV clusters at the TSS was approximately 10% lower than 5kb upstream of the TSS (Fig. 4). A similar trend was observed for the 5kb downstream regions (∼ 7%). In comparison, the absolute number of SNV around the TSS was more than ten times lower than the number of SV clusters. With the exception of a distinct peak at position two downstream of the TSS, the number of SNV around the TSS followed the same trends as described for the SV clusters above.

**Fig. 4:**
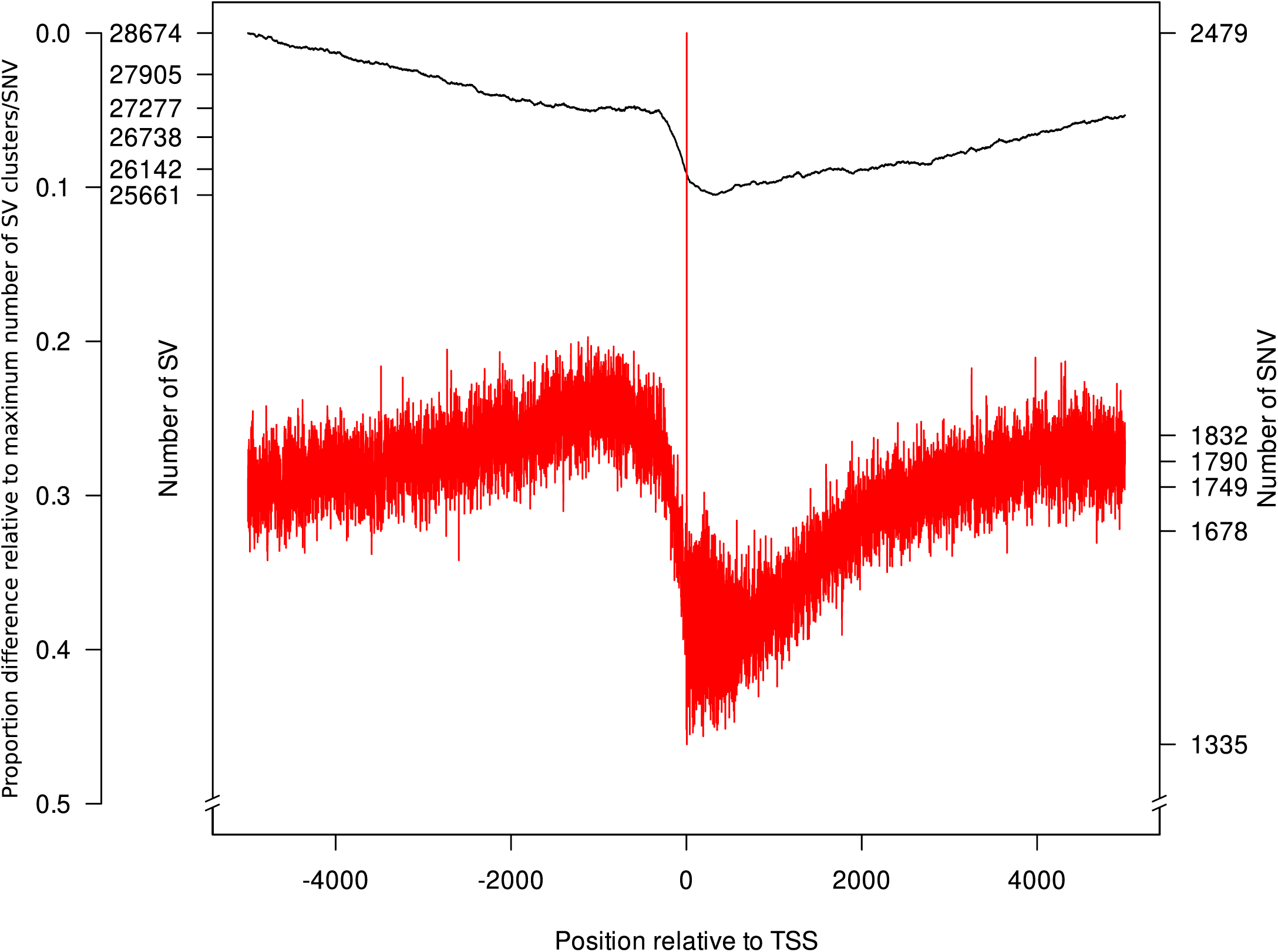
Distribution of structural variant (SV) clusters (black) and single nucleotide variants (SNV, red) among 23 barley inbreds relative to the transcription start site (TSS) of a gene (x-axis). SV clusters and SNV were counted for every position from 5kb up- and downstream around the TSS of all genes (y-axes). As third y-axis, the proportion difference relative to the maximum number of SV clusters/SNV is illustrated.

### Association of SV clusters with gene expression

We evaluated the strength of the association of the allele distribution at SV clusters with gene expression variation across the 23 inbreds. As a first step, a principal component analysis of the gene expression matrix, which included all genes and inbreds, was performed. The loadings of all 23 inbreds on principal component (PC) 1 explained 19.7% of the gene expression variation and were correlated with the presence/absence status of all inbreds for each gene-associated SV cluster. The average absolute correlation coefficient of gene-associated SV clusters and the PC1 of gene expression was 0.17 and higher than the Q_95_ of the coefficient observed for randomized presence/absence pattern and the PC1 (Supplementary Fig. S10, Supplementary Fig. S11). Similar observations were made for the association of gene-associated SV clusters with PC2 and PC3 of 0.17 and 0.19, respectively, for the above-mentioned gene expression matrix (Supplementary Fig. S12). In addition, we investigated a possible association between SV clusters and gene expression on an individual gene basis. For a total of 1,976 out of 21,140 gene-associated SV clusters a significant (P *<* 0.05) association with the gene expression of the associated gene was observed (Fig. 5).

**Fig. 5:**
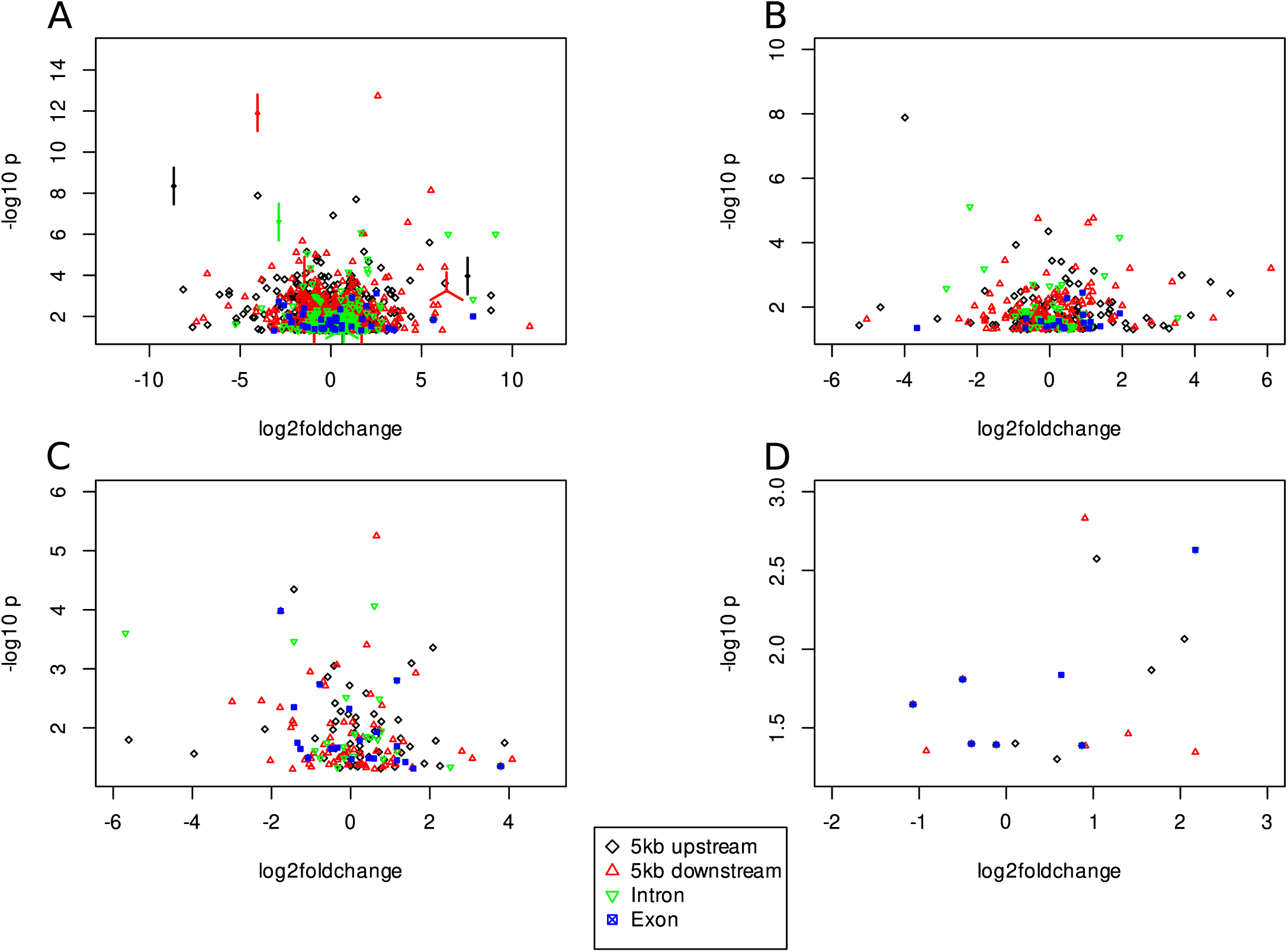
Association of gene-associated (for details see Material & Methods) deletions (A), insertions (B), duplications (C), and inversions (D) with a minor allele frequency *>* 0.15 with the expression of individual genes assessed using the PK mixed linear model. The gene-associated structural variant (SV) clusters were classified based on their occurrence relative to genes in 5kb up- or downstream, introns, and exons. Values of SV clusters with the same coordinates are illustrated as points with edges, where each edge represents one SV cluster.

### Prediction of phenotypic variation from SV clusters

The prediction ability of seven quantitative phenotypic traits using SV clusters as well as SNV from a single nucleotide polymorhpism (SNP) array, genome-wide gene expression information, SNV and INDELs (SNV&INDELs) were examined as predictors through five-fold cross-validation. The median prediction ability across all traits ranged from 0.509 to 0.648. The SV clusters had the highest prediction power, followed by SNV&INDELs, SNP array, and gene expression in decreasing order (Fig. 6). Compared to these differences, those among the median prediction abilities of the different SV types were small. The highest prediction ability was observed for insertions and the lowest for inversions. We also evaluated the possibility to combine SNV and INDELs with gene expression and SV cluster information using different weights to increase the prediction ability (Supplementary Fig. S13). The mean of the optimal weight across the seven traits was highest for gene expression (0.41) and lowest for SV clusters (0.23) (Supplementary Table S5).

**Fig. 6:**
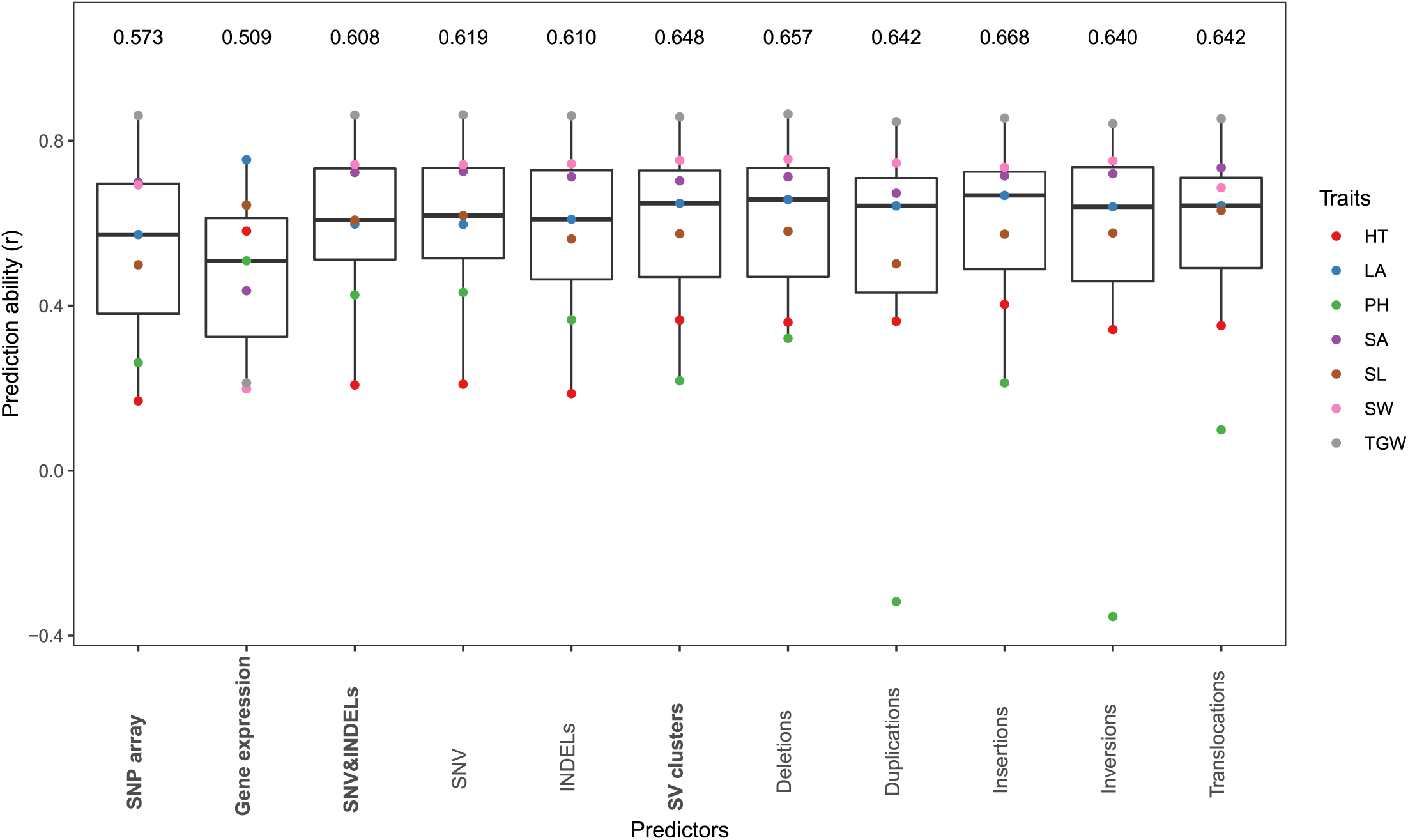
Boxplot of the median prediction abilities across the seven traits heading time (HT), leaf angle (LA), plant height (PH), seed area (SA), seed length (SL), seed width (SW), thousand grain weight (TGW) based on 23 inbreds using different predictors. The points in each box represent the medians of 200 five-fold cross-validation runs for each trait. The predictors were: features from SNP array, gene expression, single nucleotide variants (SNV) and small insertions and deletions (2 - 49bp, INDELs), as well as structural variant (SV) clusters individually as well as combined together.

## DISCUSSION

The improvements to sequencing technologies made SV detection in large genomes possible (Della Coletta et al., 2021). Despite these advances, the relative high cost of third compared to second generation sequencing makes the former less affordable and scalable for many research groups. This fact is particularly strong if genotypes have to be analyzed. We therefore used computer simulations to study the precision and sensitivity of SV detection based on different sequencing coverages of short-read sequencing data in the model cereal barley. We also evaluated whether linked-read sequencing offered by BGI (Wang et al., 2019) or formerly 10x Genomics (Weisenfeld et al., 2017) is advantageous for SV detection compared to classical Illumina sequencing.

### Precision and sensitivity to detect SV in complex cereal genomes using short-read sequencing data are high

The costs for creating linked-read sequencing libraries is considerably higher compared to that of classical Illumina libraries. Taking this cost difference into account, a fair comparison of precision and sensitivity to detect SV is between 25x Illumina and 14x linked-reads. However, even when directly compared at equal (25x) sequencing coverage, the F1-score, which is the harmonic mean of the precision and sensitivity, on average across all SV types and SV length categories was higher for Illumina compared to linked-reads (Supplementary Fig. S1). One reason might be that the SV callers used in our study do not fully exploit linked-read data. In our study, linked-read information was only used to improve the mapping against the reference genome (Marks et al., 2019). More recently, SV callers have been described that exploit linked information of linked-read data as VALOR2 (Karaoglanoglu et al., 2020) or LEVIATHAN (Morisse et al., 2021). However, the SV callers that were available at the time the simulations were performed had a very limited spectrum of SV types and SV length categories they could detect e.g. LongRanger wgs (Zheng et al., 2016) and NAIBR (Elyanow et al., 2018). In addition, we have observed for these SV callers in first pilot simulations considerably lower values for precision and sensitivity to detect SV compared to the classical short-read SV callers. Therefore, only short-read SV callers were evaluated in detail.

One further aspect that we examined was the influence of the sequencing coverage on sensitivity and precision of SV detection. Only a marginal difference between the F1-scores of the best combination of SV callers for a 25x vs. 65x Illumina sequencing coverage was observed (Supplementary Fig. S1). In addition, for some SV length categories, the F1-score for 25x compared to 65x sequencing coverage was actually higher. A possible explanation for this observation may be that a higher sequencing coverage can lead to an increased number of spuriously aligned reads (Kosugi et al., 2019). These reads can lead to an increased rate of false positive SV detection (Gong et al., 2021). Our result suggests that for homozygous genomes, Illumina short-read sequencing coverage of 25x is sufficient to detect SV with a high precision and sensitivity. We therefore made use of this sequencing coverage not only for further simulations but also to re-sequence the 23 barley inbreds of our study.

In addition, we also tested if a lower sequencing coverage could be used for SV detection to reduce the cost for sequencing further. We observed lower F-scores for all SV types using a sequencing coverage of 12.5x than for 25x (Supplementary Fig. S2). However, the F1-score was still *>* 80% for all SV types suggesting that even a sequencing coverage of 12.5x would have been suffered for SV detection in barley. When decreasing the sequencing coverage further, the precision and sensitivity to detect SV decreased considerably.

The SV callers evaluated here were chosen based on former benchmarking studies in human (Cameron et al., 2019; Chaisson et al., 2019; Kosugi et al., 2019) as well as rice (Fuentes et al., 2019) and pear (Liu et al., 2020b). Across all SV types and SV length categories, we observed the highest precision and sensitivity for Manta and GRIDSS followed by Pindel with only marginally lower values (Supplementary Table 2). This nding is in accordance with results of Cameron et al. (2019) for humans. In comparison to the results of Fuentes et al. (2019), we observed a considerably lower sensitivity and precision for Lumpy and NGSEP (Supplementary Table 2). This difference in performance of the SV callers in rice and barley might be explained by the difference in genome length as well as the high proportion of repetitive elements in the barley genome (Mascher et al., 2017).

Despite the high sensitivity and precision observed for some SV callers, we observed even higher values when using them in combination (Supplementary Table 2). This can be explained by the different detection principles such as paired-end reads, split reads, read depth, and local assembling that are underlying the different SV callers. Our observation indicates that a combined use of different short-read SV callers is highly recommended. This approach was then used for SV detection in the set of 23 spring barley inbreds.

### Validation of SV in the barley genome

A PCR based approach was used to validate a small subset of all detected SV. In accordance with earlier studies (Zhang et al., 2015; Yang et al., 2019; Guan et al., 2021), we evaluated the agreement between the detected SV and PCR results (Supplementary Fig. S3) for deletions and insertions up to 0.3kb (Supplementary Fig. S4). For eleven out of the eleven SV, we observed a perfect correspondence.

Our PCR results further suggested that the SV callers were able to detect eight out of 11 deletions between 0.3kb and 460kb (Supplementary Fig. S5) based on the short-read sequencing of the non-reference parental inbred Unumli-Arpa. In four of the eleven PCR reactions, however, more than one band was observed. This was true three times for the non-reference genotype Unumli-Arpa and one time for Morex (Supplementary Fig. S5B). In two of the four cases, PCR indicated the presence of both SV states in one genome. This was true for Morex as well as Unumli-Arpa and might be due to the complexity of the barley genome which increases the potential for off-target amplification.

In conclusion, for 19 of the 22 tested SV (Supplementary Table S2), the SV detected in the non-reference parental inbred by the SV callers was also validated by PCR. This high validation rate implies in addition to the high precision and sensitivity values observed for SV detection in the computer simulations that the SV detected in the experimental data of the 23 barley inbreds can be interpreted.

### Characteristics of SV clusters in the barley gene pool

Across the 23 spring barley inbreds that have been selected out of a world-wide diversity set to maximize phenotypic and genotypic diversity (Weisweiler et al., 2019), we have identified 458,671 SV clusters (Table 3). This corresponds to 1 SV cluster every 9,149 bp and corresponds to what was observed by Jayakodi et al. (2020). This number is in agreement with the number of SV clusters detected for cucumber (9,788 bp^−1^) (Zhang et al., 2015) or peach (8,621 bp^−1^) (Guan et al., 2021). Other studies have revealed a higher number of SV clusters than observed in our study. This might be due to the considerably higher number of re-sequenced accessions in rice (214 bp^−1^) (Fuentes et al., 2019), tomato (3,291 bp^−1^) (Alonge et al., 2020), and grapevine (1,260 bp^−1^) (Zhou et al., 2019).

The highest proportion of SV clusters detected in our study were deletions, followed in decreasing order by translocations, duplications, insertions, and inversions (Table 3). This is in disagreement with earlier studies where the frequency of duplications was considerably lower compared to that of insertions (Zhang et al., 2015; Zhou et al., 2019; Guan et al., 2021). Barley’s high proportion of duplications compared to other crops may be due to its high extent of repetitive elements (Mascher et al., 2017).

In contrast to earlier studies in grapevine and peach (e.g. Zhou et al., 2019; Guan et al., 2021) we observed a strong non-uniform distribution of SV clusters across the genome. Only 14.5% of the SV clusters were located in pericentromeric regions, which make up 25.7% of the genome, whereas the rest was located distal of the pericentromeric regions (Fig. 2). This pattern was even more pronounced for SV hotspots, i.e. regions with a significantly (P *<* 0.05) higher amount of SV clusters than expected based on the average genome-wide distribution. Almost all SV hotspots (95.5%) were located distal of the pericentromeric regions (74.3% of the genome) where higher recombination rates are observed. Our observation indicates that the majority of SV clusters in barley is caused by mutational mechanisms related to DNA recombination-, replication-, and/or repair-associated processes and is only to a low extent due to the activity of transposable elements. This is supported by the observation that, with the exception of translocations, only 1.4 to 25.2% of SV clusters were located in genome regions annotated as transposable elements (Table 3).

To complement our genome-wide analysis of barley SV clusters, we also examined their occurrence relative to genes and their association with gene expression.

### Association of SV clusters with transcript abundance

About 60% of the SV clusters were detected in the intergenic space (Fig. 3). The remaining SV clusters were gene-associated and detected in regions either 5kb up- or downstream of genes (∼30%) while ∼10% were detected in introns and exons (Fig. 3). These values are in the range of those previously reported for rice (∼75%, NA, exons: ∼6%) (Fuentes et al., 2019), potato (∼37%, ∼37%, ∼26%) (Freire et al., 2021), and peach (∼52%, ∼27%, ∼21%) (Guan et al., 2021). The higher proportion of SV clusters in genic regions in potato and peach compared to the cereal genomes might suggest that SV clusters are more frequently associated with gene expression in clonally than in sexually propagated species. A possible explanation for this observation could be the degree of heterozygosity in clonal species, which is considerably higher compared to that in selfing species such as rice and barley. Hence, it is plausible that they better tolerate SV clusters close to genes.

Our study was based on 23 barley inbreds which confer a limited statistical power to detect SV cluster-gene expression associations. However, this leads not to an increased proportion of false positive associations. Therefore, the findings are discussed here. We observed that the average absolute correlation coefficient of gene-associated SV clusters and global gene expression measured as loadings on the principal components was with 0.17 significantly (P *<* 0.05) different from 0 (Supplementary Fig. S10). In addition, 700 gene-associated SV clusters were individually associated (P *<* 0.05) with genome-wide gene expression. A further 1,976 alleles of gene-associated SV clusters were significantly (P *<* 0.05) associated with the expression of the corresponding 1,594 genes (Fig. 5). Additional support is given by the observation that despite SV clusters have a similar distribution across the genome as SNV, SV clusters covered more positions (in bp) of promoter regions than SNV (Fig. 4). These figures of significantly gene-associated SV clusters are in agreement with earlier figures for tomato (Alonge et al., 2020) and soybean (Liu et al., 2020a) and highlight the high potential of SV clusters to be associated with phenotypic traits.

### Genomic prediction

Because of the limited number of inbreds included in this study, the power to identify causal links between SV clusters and phenotypes is low when considering only the 23 inbreds. However, instead of examining the association of individual SV clusters with phenotypic traits, we evaluated their potential to predict seven phenotypic traits in comparison to various other molecular features which is expected to provide reasonable information also with a limited sample size (Weisweiler et al., 2019).

We observed that the ability to predict these seven traits was higher for SV clusters compared to the benchmark data from a SNP array (Fig. 6). This might be explained by the considerably higher number of SV clusters than variants included in the SNP array. However, we observed the same trend when comparing the prediction ability of SV clusters to that of the much more abundant SNV&INDELs. This indicates that the SV clusters comprise genetic information that is not comprised by SNV&INDELs. Our result is supported by the observation that when examining the combination of SNV and INDELs with gene expression and SV clusters to predict phenotypic traits, an increase of the prediction ability was observed compared to the ability observed for the individual predictors (Supplementary Table S5). Furthermore, our observation of a different prediction ability between SV clusters and SNV&INDELs can be explained by a lower extent of LD between SV clusters and linked SNV compared to that between SNV and linked SNV (Supplementary Table S4). These findings together illustrate the high potential of using SV clusters for the prediction of phenotypes in diverse germplasm sets. Such type of applications might be used also in commercial plant breeding programs. From a cost perspective such approaches will be realistic if SV detection is possible from low coverage sequencing. This might be possible when comprehensive reference sets of SV per species are available as was e.g. generated in our study for barley. However, this requires further research.

### Usefulness of SV information for QTL fine mapping and cloning

The inbred lines included in our study are the parents of a new resource for joint linkage and association mapping in barley, the double round robin population (HvDRR, Casale et al. 2021). This population consists of 45 biparental segregating populations with a to total of about 4,000 recombinant inbred lines and is available from the authors upon reasonable request. The detailed characterization of the SV pattern of the parental inbreds, presented in this study, will therefore be an extremely valuable information for the ongoing and future QTL fine mapping and cloning projects exploiting one or multiple of the HvDRR sub-populations.

To illustrate this, we have mapped the naked grain phenotype in six HvDRR sub-populations (HvDRR03, HvDRR04, HvDRR20, HvDRR23, HvDRR44, HvDRR46) to chromosome 7H (7H:525620758-525637446). Taketa et al. (2008) discovered a 17kb deletion harboring an ethylene response factor gene on chromosome 7H that caused naked caryopses in barley. In our study, two parental inbreds, namely Kharsila and IG128104, are naked barley. For both inbreds, the SV calls revealed the same 17kb deletion on chromosome 7H. In contrast, the deletion was absent in the 21 other parental inbreds. This illustrates the potential of exploiting SV information of parental inbreds for gene QTL and gene cloning.

## METHODS

### Benchmarking of variant callers for detecting SV and INDELs in the barley genome

#### Computer simulations

We used Mutation-Simulator (version 2.0.3) (Kühl et al., 2021) to simulate INDELs, deletions, duplications, inversions, insertions, and translocations in the first chromosome of the Morex reference sequence v2 (Monat et al., 2019) as this was the genome sequence available when our study was performed. In accordance with Fuentes et al. (2019), we considered five SV length categories for each of the above mentioned SV types (except translocations) (A: 50 - 300bp; B: 0.3 - 5kb; C: 5 - 50kb; D: 50 - 250kb; E: 0.25 - 1Mb) plus INDELs (2-49bp). Translocations were simulated for 50bp - 1Mb (ABCDE). We simulated SV with a mutation rate of 1.9x10^−6^ for the SV length categories A-C and INDELs, whereas mutation rates of 3.8x10^−6^ and 1.9x10^−7^ were assumed for SV length categories D and E, respectively. For each type of SV, we used BBMap’s randomreads.sh (BBMap - Bushnell B. - sourceforge.net/projects/bbmap/) to simulate 2x150bp Illumina reads with a sequencing coverage of 1.5x, 3x, 6x, 12.5x, 25x, and 65x as well as LRSim (version 1.0) (Luo et al., 2017) to simulate linked-reads with a sequencing coverage of 14x and 25x. Illumina- and linked-reads were simulated with a minimum, average, and maximum base quality of 25, 35, and 40, respectively.

#### SV detection

The simulated Illumina reads were mapped to the first chromosome of the Morex reference sequence v2 using BWA-MEM (version 0.7.15) whereas LongRanger align (version 2.2.2) was used for the simulated linked-reads. The SV callers Pindel (version 0.2.5b9) (Ye et al., 2009), Delly (version 0.8.1) (Rausch et al., 2012), GRIDSS (version 2.8.3) (Cameron et al., 2017), Manta (version 1.6.0) (Chen et al., 2016), Lumpy (smoove version 0.2.5) (Layer et al., 2014), and NGSEP (version 3.3.2) (Duitama et al., 2014) were used to identify SV based on the mapped reads. GATK’s HaplotypeCaller (4.1.6.0) (Poplin et al., 2017), Pindel, and GRIDSS were used to detect INDELs. The work ow was implemented in Snakemake (version 5.10.0) (Köster et al., 2021). A SV call was only kept if it passed the built-in filter of the corresponding SV caller. We calculated the sensitivity (1), precision (2), and the F1-score (3) as

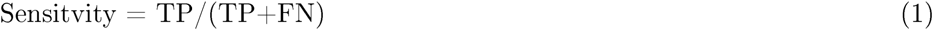

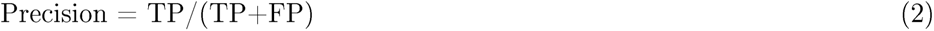

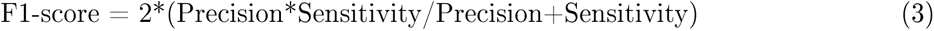

for all combinations of SV types*SV callers, where TP was the number of true positives, FP the number of false positives, and FN the number of false negatives. For INDELs, a TP INDEL had break points that did differ ≤ 2bp from those of the simulated INDEL and the length did differ by ≤ 5bp. For SV length category A, a TP SV had break points that did differ ≤ 10bp from those of the simulated SV and the SV length did differ by ≤ 20bp. For the other SV length categories, a TP SV had break points and length differences compared to the simulated SV of ≤ 50bp. For insertions where no SV length was detected, the start of a TP insertion had a break point that did differ ≤ 10bp from this of the simulated insertion. For translocations, a TP translocation had break points that did differ ≤ 50bp from those of the simulated translocation.

We also evaluated combinations of SV callers for their precision and sensitivity to detect SV. The following procedure was used to decide for the combinations that were examined: First, for those SV callers, which have shown a precision ≥ 95% for all SV length categories for a particular SV type, SV calls were combined via logical or (“|”). Second, for those SV callers with a precision ≤ 95% in at least one SV length category, SV calls were combined with a logical and (“&”). If the precision of the combinations of the second step increased to ≥ 95% in all SV length categories, SV calls of this combinations were kept for the particular SV type and were combined with a logical or with those of the first step.

The threshold of ≥ 95% precision was used to reduce the number of FP SV calls to a reasonable level.

### Detection of SV, SNV, and INDELs in the barley genome

#### Genetic material and sequencing

Our study was based on 23 spring barley inbreds (Weisweiler et al., 2019) that were selected out of a worldwide collection of 224 inbreds (Haseneyer et al., 2010) (Supplementary Table S6) using the MSTRAT algorithm (Gouesnard, 2001). These inbreds are the parents of the double round robin population (Casale et al. 2021). Paired-end sequencing libraries with an insert size of 425bp were sequenced to a ∼25x coverage on the Illumina HiSeqX platform by Novogene Corporation Inc. (Sacramento, USA).

#### SV, INDELs, and SNV detection

The quality of the raw reads was checked by fastqc. Reads were adapter- and quality-trimmed using Trimmomatic (version 0.39) (Bolger et al., 2014). The trimmed reads were mapped to the Morex reference sequence v3 (Mascher et al., 2021) using BWA-MEM. PCR-duplicates were removed using PICARD (version 2.22.0).

Based on the results of the benchmarking of different SV callers using simulated data, results of specific SV callers were combined as explained above. The final set of deletions for each inbred were those that were identified by Manta | GRIDSS | Pindel | Delly | (Lumpy & NGSEP) where homozygous-reference (0/0) and heterozygous allele (0/1) calls were removed. Additionally, deletions annotated as “replacement” (RPL) by Pindel were removed. In analogy, the duplications were identified by Manta | GRIDSS | Pindel | (Delly & Lumpy). Insertions of the SV length category A were identified by Manta | GRIDSS | Delly, where insertions of the SV length categories B-E were called using Manta. Inversions were identified by Manta | GRIDSS | Pindel. Translocations were called from pairs of break points identified by Manta | GRIDSS | (Delly & Lumpy). INDELs were detected by GATK’s HaplotypeCaller | GRIDSS | Pindel. SV which were located in a region of the reference sequence, where the sequence only consists of N’s, were excluded. For genome regions, where break points of different SV overlapped or were inconsistent in the same inbred, only the smallest SV was considered. The SV of the 23 inbreds were grouped together to SV clusters based on the similarity of sizes and the position in the genome according to the following procedure. The distance from a SV to the next SV in such a SV cluster had to be smaller than 20bp for the SV length category A and 50bp for the SV length category B - E and the difference of the two break points had to be smaller than 10 or 50bp as described above. SV with a larger difference between break points were kept as separate SV and SV clustering was pursuing. Each SV cluster was genotyped across the examined 23 barley inbreds.

SNV and INDELs were called using GATK. First, GATK’s HaplotypeCaller was used in single sample GVCF mode, afterwards GATK’s CombineGVCFs was used to combine the SNV across the 23 inbreds. Combined SNV were genotyped using GATK’s GenotypeGVCFs. SNV were ltered using GATK’s VariantFiltration (QD *<* 2.0; QUAL *<* 30.0; SOR *>* 3.0; FS *>* 60.0; MQ *<* 40.0; MQRankSum *<* -12.5; ReadPosRankSum *<* -8.0).

#### PCR validation of SV

A total of 25 of the detected SV were targeted for validation by PCR amplification of genome regions of and around the SV in Morex and Unumli-Arpa. This included six SV length category A deletions, five SV length category A insertions, six SV length category B deletions and eight SV length category C-E deletions. In order to determine the SV allele, we required the amplification of two differently sized fragments in the two inbreds. For each SV, a regular primer pair was created with the position defined by the validation strategy (Supplementary Fig. S1). If needed, a second right primer was added to the PCR reaction. The primers were designed using Primer3 (Untergasser et al., 2012) and Blast+ (Camacho et al., 2009).

Plant material was sampled for the PCR validation from adult plants and seedlings grown under controlled conditions. DNA was extracted from 100 mg frozen plant material using the DNeasy Plant Mini Kit (Qiagen, Germany) according to the manufacturer’s instructions. The PCR reaction mixture contained in a nal volume of 20 *µ*L: 0.2 mM dNTP, Fw/Rev Primer 0.5 *µ*M, 50 ng DNA, 1.5 U/*µ*L DreamTaq DNA Polymerase (Thermo Fischer Scientific, USA), Polymerase-Buffer 1X and water. Amplified fragments were separated by gel electrophoresis and the validation success was determined by comparing the PCR product sizes with the calculated values based on the SV detection.

#### Location of SV clusters

SV clusters were classified and annotated based on their location in the genome, their distance relative to genes, or other genomic features. SV clusters were grouped into four gene-associated and one intergenic SV cluster categories: 5kb upstream/downstream gene-associated SV clusters were located in the 5kb region from the 3′- or 5′- end of a gene. Intron and exon gene-associated SV clusters were located in the gene sequence, where the genic sequence was separated into intronic and exonic sequences. SV clusters which were not located in the four gene-associated SV cluster categories were determined as intergenic SV clusters. A gene-associated SV cluster could be classified in more than one category if its sequence covers several genomic features.

To check if the detected SV clusters were transposable elements, the genomic positions of SV clusters were compared to the transposable elements annotation le of the Morex reference sequence v3 (Mascher et al., 2021). Deletions, duplications, inversions, INDELs, and insertions with known length were annotated as transposable elements if the reciprocal overlap was ≥ 80% (Fuentes et al., 2019). Insertions with unknown length were classified as transposable elements if the detected break point of the insertion was inside the transposable element sequence. Translocations were classified as transposable element, if at least one of the two break points was located inside a transposable element sequence.

SV hotspots were identified using the following procedure: The average number of SV clusters in non-overlapping 1Mb windows across each of the seven chromosomes was determined. Using this number, we calculated for each window based on the poisson distribution the expected number of SV clusters. Windows with more SV clusters than the Q_99_ of the expected poisson distribution were designated as SV hotspots (Guan et al., 2021).

#### Population genetic analyses

LD measured as r^2^ (Hill and Robertson, 1968) was calculated between each SV type and linked SNV. Nucleotide diversity (*π*) was calculated in 100kb windows along the seven chromosomes separately for SV clusters (deletions, insertions, duplications, inversions) and SNV using vcftools (version 0.1.17) (Danecek et al., 2011).

#### SV clusters and gene expression

SV clusters which were assigned into one of the gene-associated SV categories, namely 5kb up- or downstream, introns, and exons, were associated with the genome-wide gene expression of the 23 barley inbreds. Gene expression for the seedling tissue measured as fragments per kilobase of exon model per million fragments mapped was available for all inbreds from an earlier study (Weisweiler et al., 2019). This information was the basis of a principal component analysis. For all gene-associated SV clusters with a MAF *>* 0.15, Pearson’s correlation coefficient with the first three principal components was estimated, where presence and absence of SV clusters were used as metric character. A permutation procedure with 1,000 iterations was used to test the mean absolute values of the correlations for their significance. In addition to this evaluation of the effect of SV clusters on the genome-wide gene expression level, we also examined the significance of the effect of gene-associated SV clusters with a MAF *>* 0.15 on the expression of individual genes. In order to do so, the mixed linear model with population structure and kinship matrix (PK model) (Stich et al., 2008) was used. The population structure matrix consisted of the first two principal components calculated from 133,566 SNV and INDELs derived from mRNA sequencing (Weisweiler et al., 2019). From the same information, the kinship matrix was calculated as described by Endelman and Jannink (2012).

#### Assessment of phenotypic traits

For the assessment of phenotypic traits under field conditions, the 23 inbreds were planted as replicated checks in an experiment laid out as an augmented row-column design. The experiment was performed in seven agro-ecologically diverse environments (Cologne from 2017 to 2019, Mechernich and Quedlinburg from 2018 to 2019) in Germany in which the checks were replicated multiple times per environment. For each environment, seven phenotypic traits were assessed. Heading time (HT) was recorded as days after planting, leaf angle (LA) was scored on a scale from 1 (erect) to 9 (very flat) on four-week-old plants, and plant height (PH, cm) was measured after heading in Cologne and Mechernich. Seed area (SA, mm^2^), seed length (SL, mm), seed width (SW, mm), and thousand grain weight (TGW, g) were measured based on full-filled grains from Cologne (2017-2019) and Quedlinburg (2018) by using MARVIN seed analyzer (GTA Sensorik, Neubrandenburg, Germany).

#### Prediction of phenotypes

Each of the phenotypic traits was analyzed across the environments using the following mixed model:

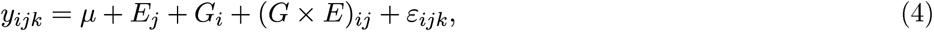

where *y_ijk_* was the observed phenotypic value for the *i^th^* genotype at the *j^th^* environment within the *k^th^* replication; *µ* the general mean, *G_i_*the effect of the *i^th^*inbred, *E_j_* the effect of the *j^th^* environment, (*G* × *E*)*_ij_*the interaction between the *i^th^*inbred and the *j^th^* environment, and *ε_ijk_* the random error. This allowed to estimate adjusted entry means for all inbreds.

The performance to predict the adjusted entry means of each barley inbred for each trait using different types of predictors: (1) SNP array, which was generated by genotyping the 23 inbreds using the Illumina 50K barley SNP array (Bayer et al., 2017), (2) gene expression (3) SNV&INDELs, (3a) SNV, (3b) INDELs, (4) SV clusters, (4a) deletions, (4b) duplications, (4c) insertions, (4d) inversions, (4e) translocations, was compared based on genomic best linear unbiased prediction (GBLUP) (VanRaden, 2008).

For each predictor, the monomorphic features and the features with missing rates *>* 0.2 and identical information were discarded. **W** was defined as a matrix of feature measurement for the respective predictor. The dimensions of **W** were the number of barley inbreds (*n* = 23) times the number of features in the corresponding predictor (m) (*m_SNP array_* = 38, 025, *m_gene expression_* = 67, 844, *m_SNV_* _&*INDELs*_ = 3, 025, 217, *m_SNV_*= 2, 338, 565, *m_INDELs_* = 686, 652, *m_SV clusters_* = 458, 330, *m_deletions_*= 183, 219, *m_duplications_* = 93, 073, *m_insertions_* = 70, 143, *m_inversions_* = 6, 582, *m_translocations_* = 105, 313). The additive relationship matrix **G** was defined as 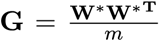 , where **W**^∗^ was a matrix of feature measurement for the respective predictor, whose columns are centered and standardized to unit variance of **W**, and **W**^∗^**^T^** was the transpose of **W**^∗^.

Furthermore, to investigate the performance of a joined weighted relationship matrix (Schrag et al., 2018) to predict phenotypic variation, the three **G** matrices in GBLUP model of the three predictors, SNV&INDELs, gene expression, and SV clusters, were weighted and summed up to one joined weighted relationship matrix. A grid search, varying any weight (*w*) from 0 to 1 in increments of 0.1, resulted in 66 different combinations of joined weighted relationship matrix, where the summation of three weights in each combination must be equal to 1.

Five-fold cross-validation was used to assess the model performance. Prediction abilities were obtained by calculating Pearson’s correlations between observed (*y*) and predicted (*y*^) adjusted entry means in the validation set of each fold. The median prediction ability across the five folds within each replicate was calculated and the median of the median across the 200 replicates was used for further analyses.

## DECLARATIONS

### Availability of data and materials

Raw DNA sequencing data of the 23 barley inbreds have been deposited into the NCBI Sequence Read Archive (SRA) under the accession PRJNA77700 and will become available after manuscript acceptance (https://dataview.ncbi.nlm.nih. gov/object/PRJNA777004?reviewer=el83fbl241mgqmbjdireuafcic). Raw mRNA sequencing data are available under the accession PRJNA534414. Data of gene expression, SNP array, adjusted entry means of phenotypes, INDELs, and SV will become available after manuscript acceptance via gshare (https://doi.org/10.6084/m9.figshare.16802473). SNV data will become available after manuscript acceptance via zenodo (https://doi.org/10.5281/zenodo.6451025). Snakemake work ows are available via github (https://github.com/mw-qggp/SV_barley). Further scripts are available from the authors upon request.

## Acknowledgements

Computational infrastructure and support were provided by the Center for Information and Media Technology (ZIM) at Heinrich Heine University D sseldorf.

## Authors’ contributions

MW and BS designed and coordinated the project; TH extracted DNA and prepared the libraries; DVI contributed phenotypic data; MW, CA, and PW performed the analyses; MW and BS wrote the manuscript.

## Ethics approval and consent to participate

The authors declare that the experimental research on plants described in this paper complied with institutional and national guidelines.

## Consent for publication

All authors read and approved the final manuscript.

## SUPPLEMENTARY INFORMATION

**Table S1:** Sensitivity/precision of structural variant (SV) callers and combinations of them (for details see Material & Methods) to identify small insertions and deletions (2 - 49bp, INDELs) and translocations (50bp - 1Mb).

| SV caller | Deletions (2 - 49bp) | Insertions (2 - 49bp) | Translocations (50bp - 1Mb) |
| --- | --- | --- | --- |
| Delly |  |  | 85.6/76.0 |
| Manta |  |  | 89.4/100.0 |
| Lumpy |  |  | 83.2/82.4 |
| GRIDSS | 68.0/99.3 | 64.6/98.9 | 87.2/100.0 |
| Pindel | 92.4/97.9 | 87.5/98.7 |  |
| GATK | 92.3/97.6 | 94.6/98.7 |  |
| Combination | 95.5/98.9 | 94.8/98.7 | 95.4/99.8 |

**Table S2:** Predicted structural variants (SV) for PCR validation. Listed are all SV that were PCR validated including the names, sizes, primer positions, and the expected amplicon sizes. All sizes are given in bp.

| SV names | Primer position relative to SV start |  |  | SV size | Expected amplicon size |  |
| --- | --- | --- | --- | --- | --- | --- |
|  | left | right | 2nd right |  | Morex | Unumli-Arpa |
| Del_A_1 | -263 | 321 |  | 57 | 584 | 527 |
| Del_A_2 | -158 | 293 |  | 64 | 451 | 387 |
| Del_A_3 | -110 | 324 |  | 53 | 434 | 381 |
| Del_A_4 | -229 | 424 |  | 124 | 653 | 529 |
| Del_A_5 | -216 | 265 |  | 55 | 481 | 426 |
| Del_A_6 | -277 | 155 |  | 59 | 432 | 373 |
| Ins_A_1 | -167 | 243 |  | 57 | 353 | 410 |
| Ins_A_2 | -238 | 191 |  | 76 | 353 | 429 |
| Ins_A_3 | -234 | 258 |  | 91 | 401 | 492 |
| Ins_A_4 | -288 | 126 |  | 52 | 362 | 414 |
| Ins_A_5 | -266 | 239 |  | 57 | 448 | 505 |
| Del_B_1 | -391 | 2,704 |  | 1,937 | 3,095 | 1,158 |
| Del_B_2 | -462 | 4,446 |  | 4,144 | 4,908 | 764 |
| Del_B_3 | -374 | 3,687 |  | 2,940 | 4,061 | 1,121 |
| Del_C_1 | -364 | 316 | 11,313 | 10,778 | 680 | 899 |
| Del_C_2 | -103 | 280 | 5,692 | 5,355 | 383 | 440 |
| Del_C_3 | -231 | 375 | 28,406 | 27,937 | 606 | 700 |
| Del_D_1 | -262 | 120 | 287,036 | 286,558 | 382 | 740 |
| Del_D_2 | -361 | 371 | 91,956 | 91,411 | 732 | 906 |
| Del_D_3 | -248 | 224 | 54,918 | 54,481 | 472 | 685 |
| Del_E_1 | -169 | 348 | 460,621 | 460,240 | 517 | 550 |
| Del_E_2 | -279 | 239 | 405,578 | 405,029 | 518 | 828 |

**Table S3:** Proportion (%) of SV length categories for deletions, duplications, inversions, and insertions.

| SV length category | Deletions | Duplications | Inversions | Insertions |
| --- | --- | --- | --- | --- |
| A (50 - 300bp) | 41.7 | 16.2 | 20.1 | 48.4 |
| B (0.3 - 5kb) | 30.3 | 21.7 | 16.5 | 5.7<sup>1</sup> |
| C (5 - 50kb) | 26.4 | 55.9 | 25.9 |  |
| D (50 - 250kb) | 1.5 | 5.5 | 24.4 |  |
| E (0.25 - 1 Mb) | 0.1 | 0.7 | 13.1 |  |

<sup>1</sup>0.3 - 1kb; no insertion length detected for 45.9%

**Table S4:**
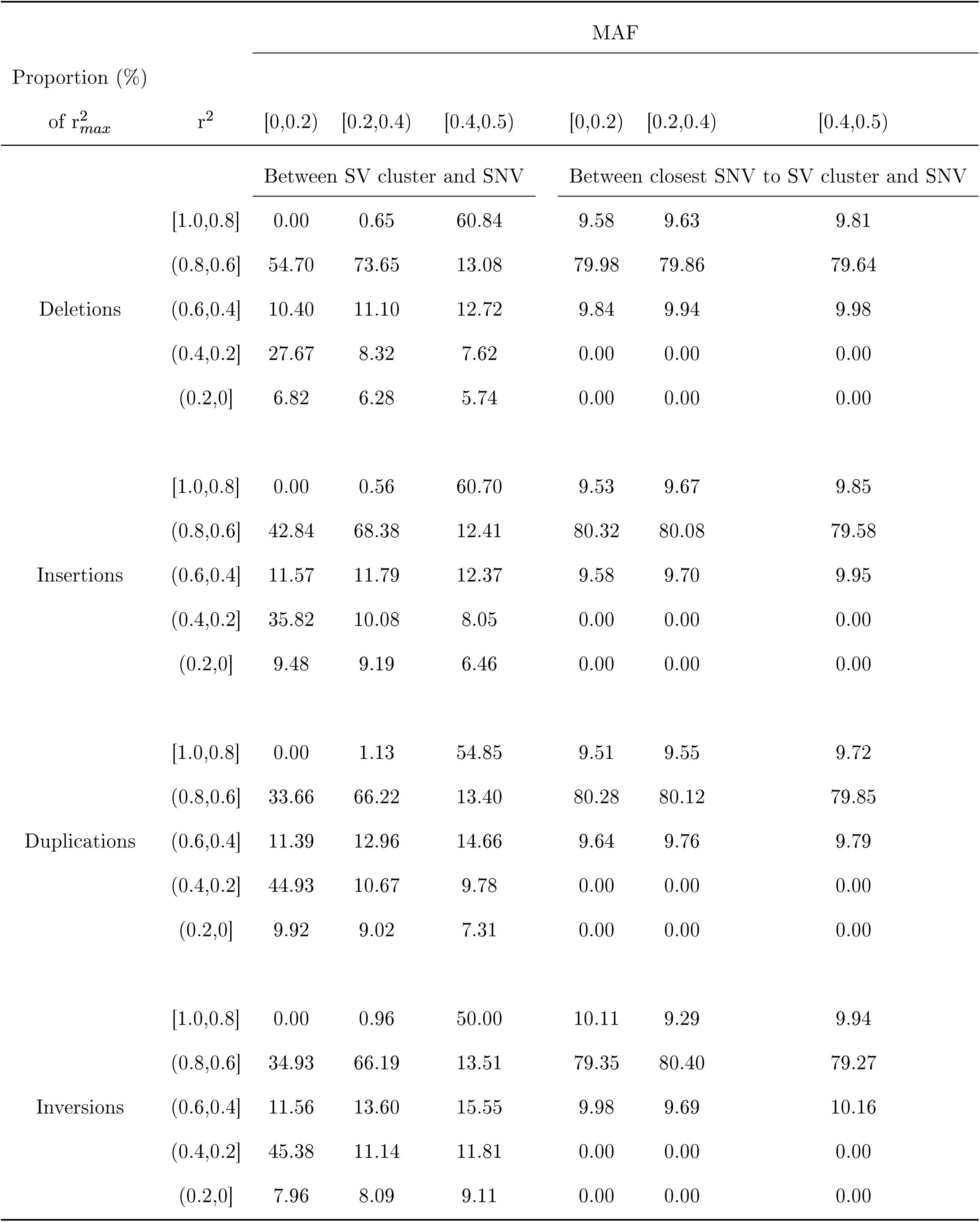
Percentage of structural variant (SV) clusters or their closest neighboring single nucleotide variant (SNV) that show a maximum linkage disequilibrium (LD) estimate r^2^*_max_* to all SNV 1kb up and downstream of it. LD was calculated for three categories of minor allele frequencies (MAF) for SV clusters and the corresponding closest SNV.

**Table S5:** The optimal weights of the three predictors single nucleotide variants (SNV) and Indel (SNV&Indel), structural variants (SV) and gene expression that resulted in the highest prediction abilities for the seven traits heading time (HT), leaf angle (LA), plant height (PH), seed area (SA), seed length (SL), seed width (SW), and thousand grain weight (TGW).

| Traits | SNV&INDELs | SV clusters | Gene expression | Prediction ability |
| --- | --- | --- | --- | --- |
| HT | 0.0 | 0.1 | 0.9 | 0.63 |
| LA | 0.0 | 0.4 | 0.6 | 0.79 |
| PH | 0.0 | 0.1 | 0.9 | 0.54 |
| SA | 0.9 | 0.0 | 0.1 | 0.74 |
| SL | 0.6 | 0.0 | 0.4 | 0.70 |
| SW | 0.0 | 1.0 | 0.0 | 0.75 |
| TGW | 1.0 | 0.0 | 0.0 | 0.86 |
| Mean (median) | 0.36 (0) | 0.23 (0.1) | 0.41 (0.4) |  |

**Table S6:** Inbred lines included in this study, their country of origin (CoO), row type, and year of release.

| Inbred name | BCC code | CoO | Row type | Year of release | Genome sequencing coverage |  |  |
| --- | --- | --- | --- | --- | --- | --- | --- |
|  |  |  |  |  | seq | seq-trimmed | mapped |
| HOR1842 | HOR1842 | AFG | 6 | 1935 | 27.4 | 26.3 | 25.9 |
| HOR383 | BCC1561 | BGR | 6 | unknown | 24.8 | 23.8 | 22.4 |
| Sanalta | BCC929 | CAN | 2 | 1930 | 27.5 | 26.3 | 25.5 |
| ItuNative | BCC502 | CHN | 6 | unknown | 23.6 | 22.7 | 21.3 |
| Sissy | BCC1413 | GER | 2 | 1990 | 24.0 | 23.1 | 22.7 |
| Georgie | BCC1381 | GBR | 2 | 1975 | 25.1 | 24.1 | 23.7 |
| SprattArcher | BCC1415 | GBR | 2 | 1943 | 23.1 | 22.2 | 22.4 |
| Lakhan | BCC533 | IND | 6 | unknown | 21.6 | 20.8 | 20.1 |
| Kharsila | HOR11403 | IND | 6 | before 1911 | 26.7 | 25.6 | 24.2 |
| W23829/803911 | HOR11374 | ISR | 2 | unknown | 23.6 | 22.7 | 22.4 |
| Namhaebori | BCC667 | KOR | 6 | unknown | 22.3 | 20.4 | 21.6 |
| IG128216 | BCC118 | LBY | 6 | 1983 | 21.2 | 19.3 | 20.8 |
| IG128104 | BCC173 | PAK | 6 | 1974 | 23.8 | 22.9 | 22.4 |
| K10693 | BCC1491 | RUS | 6 | unknown | 21.0 | 20.2 | 19.8 |
| IG31424 | BCC190 | SYR | 2 | 1981 | 23.5 | 22.5 | 21.9 |
| HOR12830 | HOR12830 | SYR | 6 | unknown | 25.8 | 24.7 | 23.4 |
| HOR7985 | HOR7985 | TUR | 2 | before 1969 | 23.3 | 22.3 | 22.3 |
| K10877 | BCC1503 | TKM | 6 | unknown | 25.5 | 24.4 | 23.7 |
| HOR8160 | HOR8160 | TUR | 2 | before 1969 | 24.4 | 23.5 | 23.0 |
| Ancap2 | BCC807 | URY | 6 | 1950 | 27.0 | 25.9 | 24.6 |
| CM67 | BCC846 | USA | 6 | 1983 | 23.8 | 22.9 | 22.3 |
| Kombyne | BCC893 | USA | 6 | 1975 | 21.5 | 20.5 | 19.9 |
| Unumli-Arpa | BCC1470 | UZB | 2 | unknown | 23.5 | 22.6 | 22.1 |

**Fig. S1:**
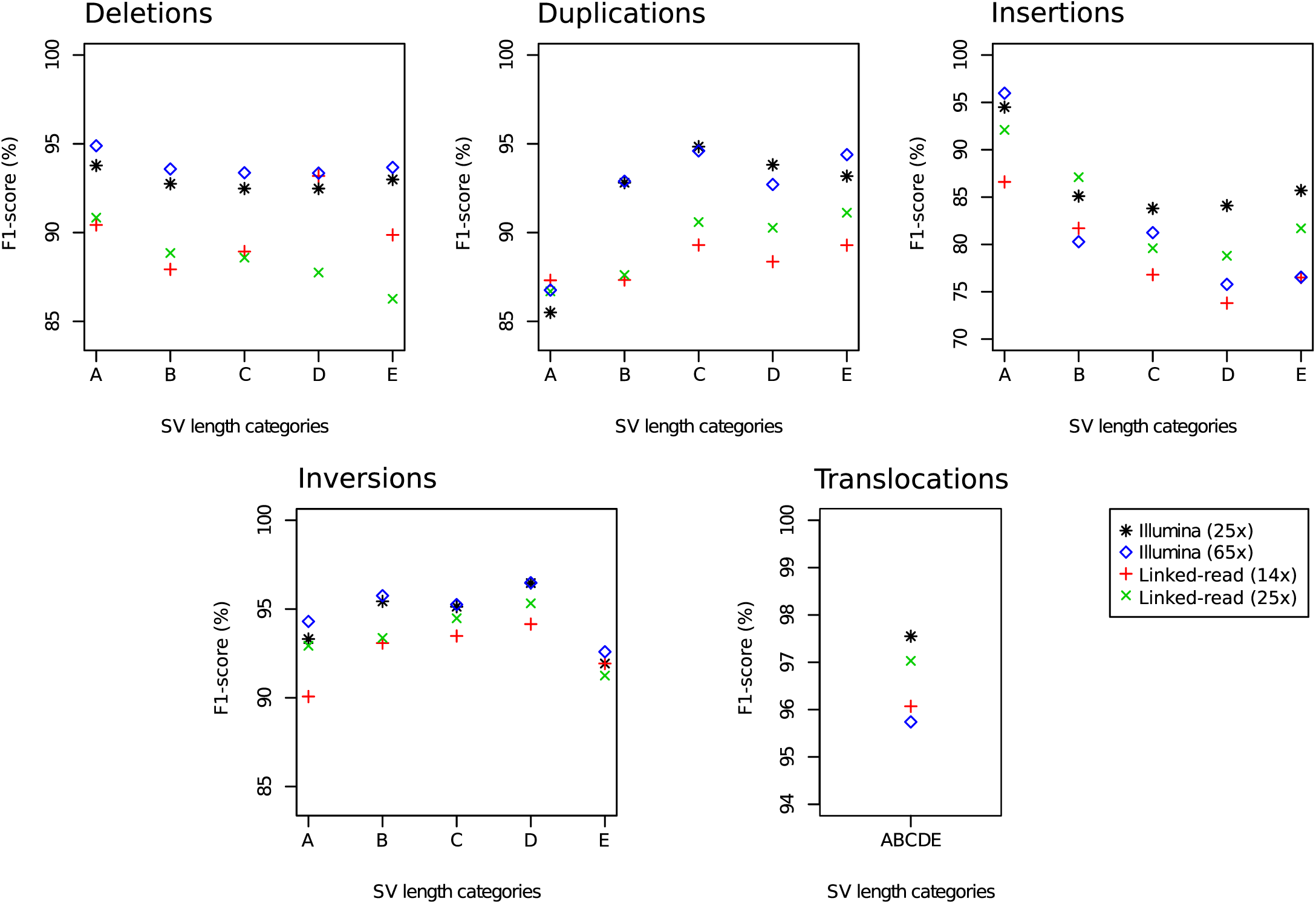
F1-score, which is the harmonic mean of the precision and sensitivity, for the detection of deletions, duplications, insertions, inversions, and translocations of five structural variant (SV) length categories: A (50 - 300bp), B (0.3 - 5kb), C (5 - 50kb), D (50 - 250kb), E (0.25 - 1Mb) using the best combination of SV callers (for details see Material & Methods) based on 25x and 65x Illumina short-read sequencing as well as based on 14x and 25x linked-read sequencing coverage.

**Fig. S2:**
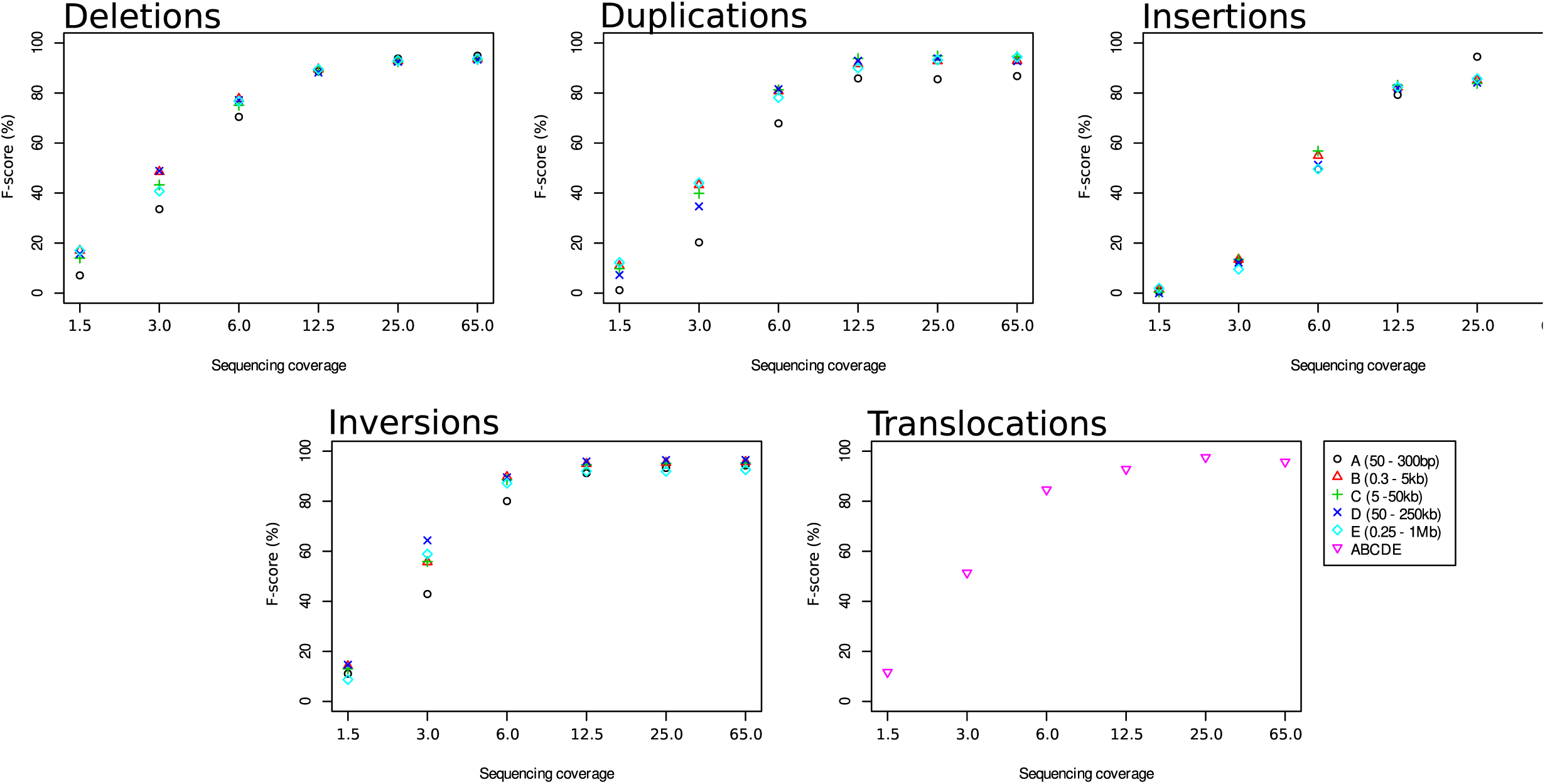
F1-score, which is the harmonic mean of the precision and sensitivity, for the detection of deletions, duplications, insertions, inversions, and translocations of six sequencing coverages (1.5x, 3.0x, 6.0x, 12.5x, 25.0x, and 65.0x) using the best combination of SV callers (for details see Material & Methods).

**Fig. S3:**
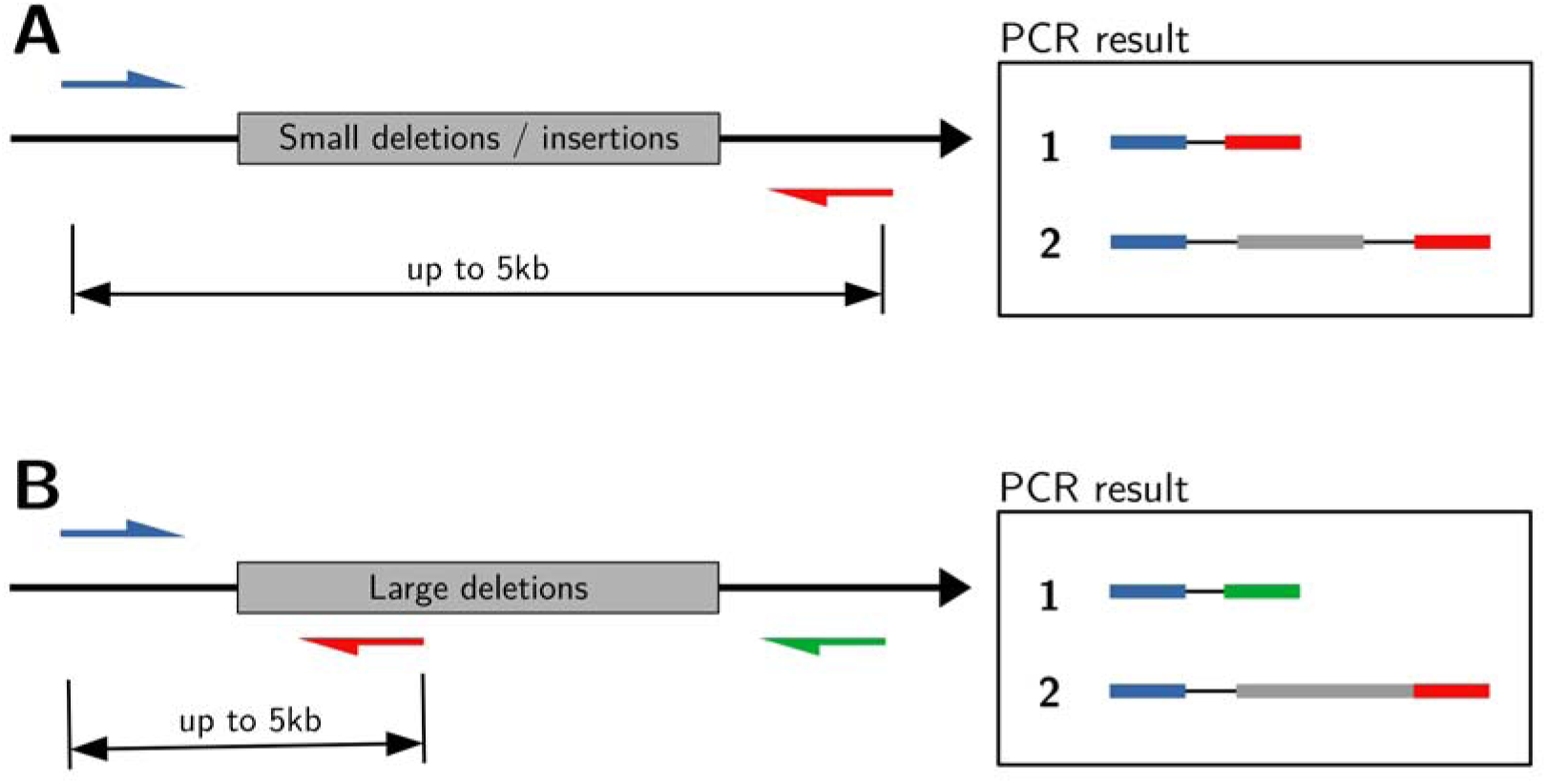
Graphical illustration of the primer design strategy created to validate structural variant (SV) predictions in the reference genome Morex and Unumli-Arpa. The primer design strategy had to be adjusted depending on the size of the SV. Smaller deletions (A) and insertions (up to ∼5kb) were validated with a pair of two primers (blue/red arrow) anking the SV (gray box). Larger deletions (B) were validated either by primer 1 (blue) and primer 2 (red) in case of presence or by primer 1 (blue) and primer 3 (green) in case of absence. The predicted PCR results, the absence (1) and presence (2) of the SV sequence in the PCR fragment, are shown on the right.

**Fig. S4:**
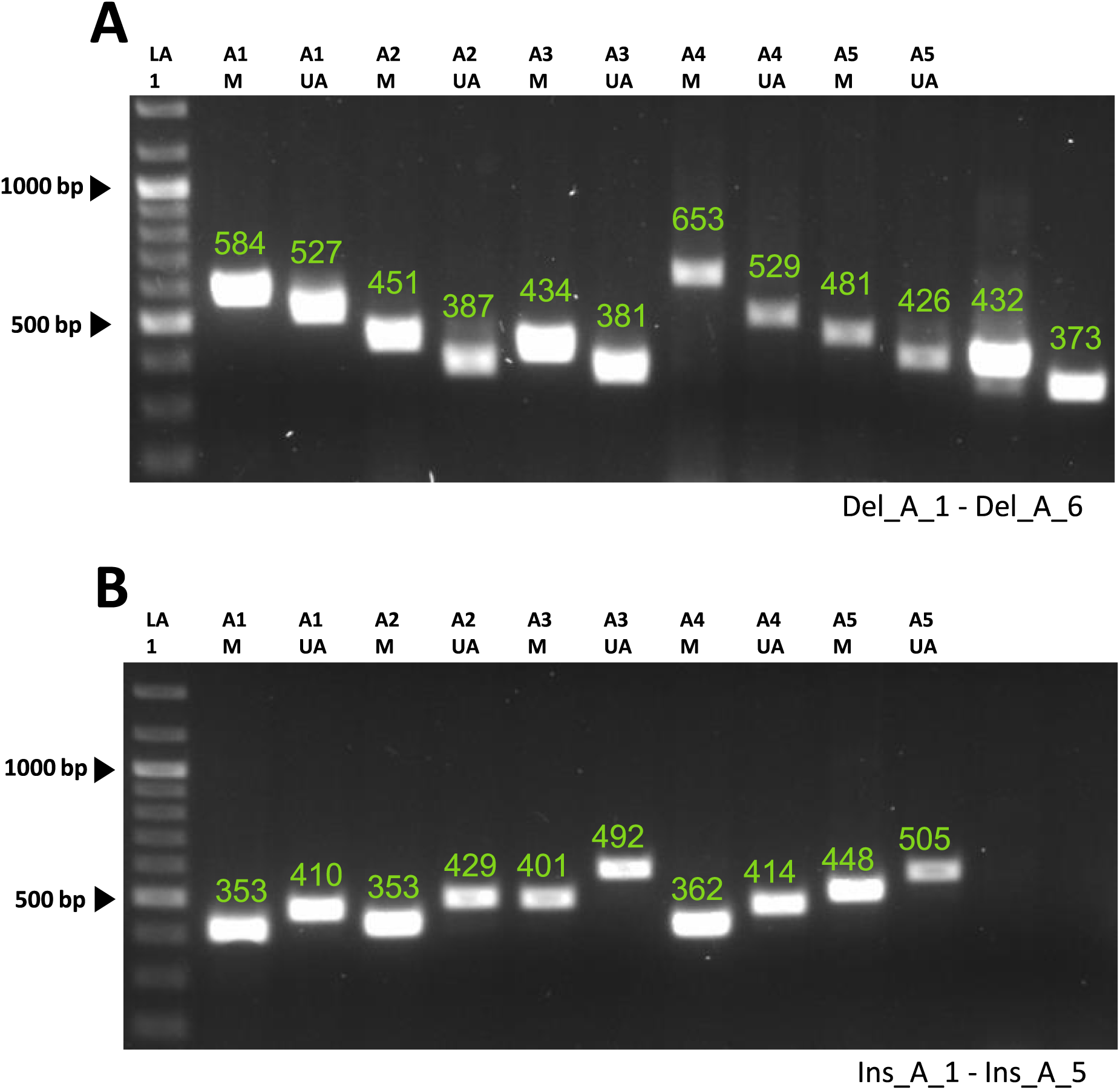
PCR validation results for small structural variants (SV) as documented after the gel electrophoresis. PCR amplified fragments are shown separated by size for the reference genotype Morex (M) and the genotype Unumli-Arpa (UA). Predicted fragment size based on the SV predictions are illustrated by numbers. The numbers are colored based on the validation success. Fragment size agreement between PCR and prediction (green) or disagreement (red). Results are shown for six small deletions (A) and six small insertions (B) of the SV length category A (50 - 300bp). DNA ladder used: GeneRuler 100bp Plus, Thermo Fisher (LA 1).

**Fig. S5:**
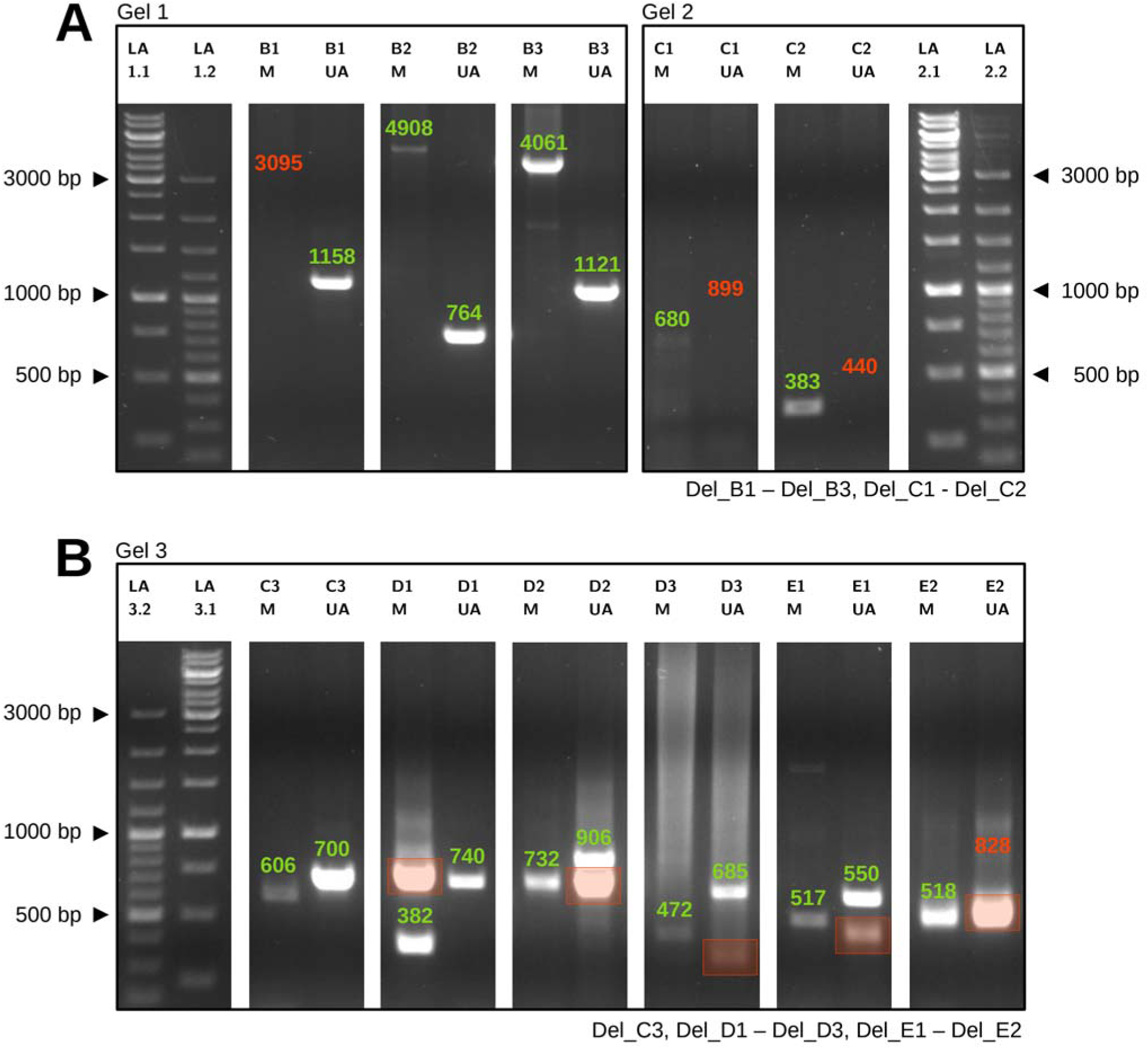
PCR validation results for large structural variants (SV) as documented after the gel electrophoresis. PCR amplified fragments are shown separated by size for the reference genotype Morex (M) and the genotype Unumli-Arpa (UA). Predicted fragment size based on the SV predictions are illustrated by numbers. The numbers are colored based on the validation success. Fragment size agreement between PCR and prediction (green) or disagreement (red). Additional not predicted fragments are marked by a red box. Results are shown for six deletions of the SV length category B (0.3 - 5kb) (A) and 8 deletions of the SV length category C (5 - 50kb), D (50 - 250kb), and E (0.25 - 1Mb) (B). DNA ladder used: GeneRuler 100bp Plus (LA 1) and GeneRuler 1kb, Thermo Fisher (LA 2).

**Fig. S6:**
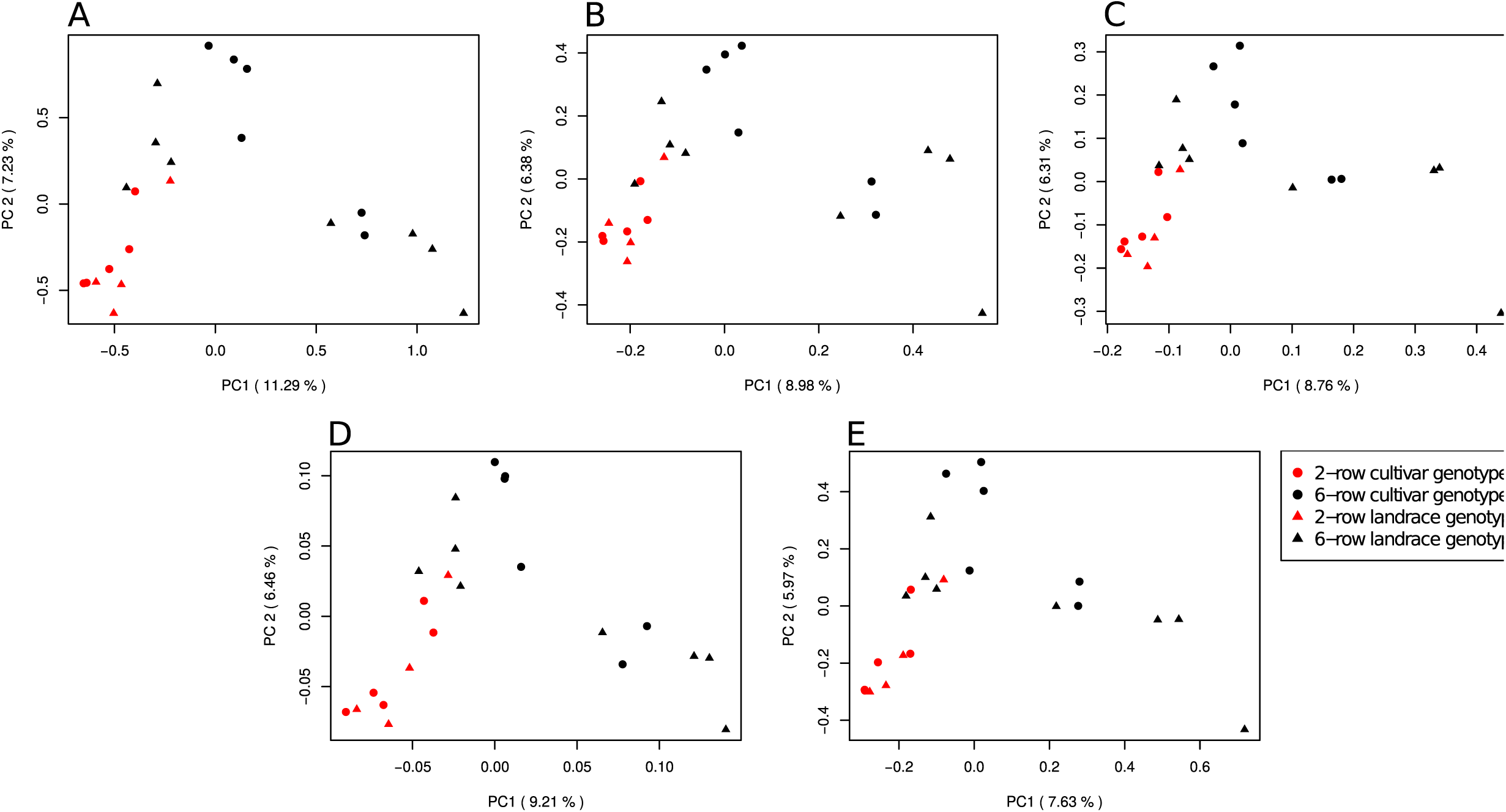
Principal component analyses of the barley inbred lines considered in our study based on deletions (A), duplicatons (B), insertions (C), inversions (D), and translocations (E). PC 1 and PC 2 are the first and second principal component, respectively, and number in parentheses refer to the proportion of variance explained by the principal components. Symbols identify landrace and cultivar inbreds and colors their row number.

**Fig. S7:**
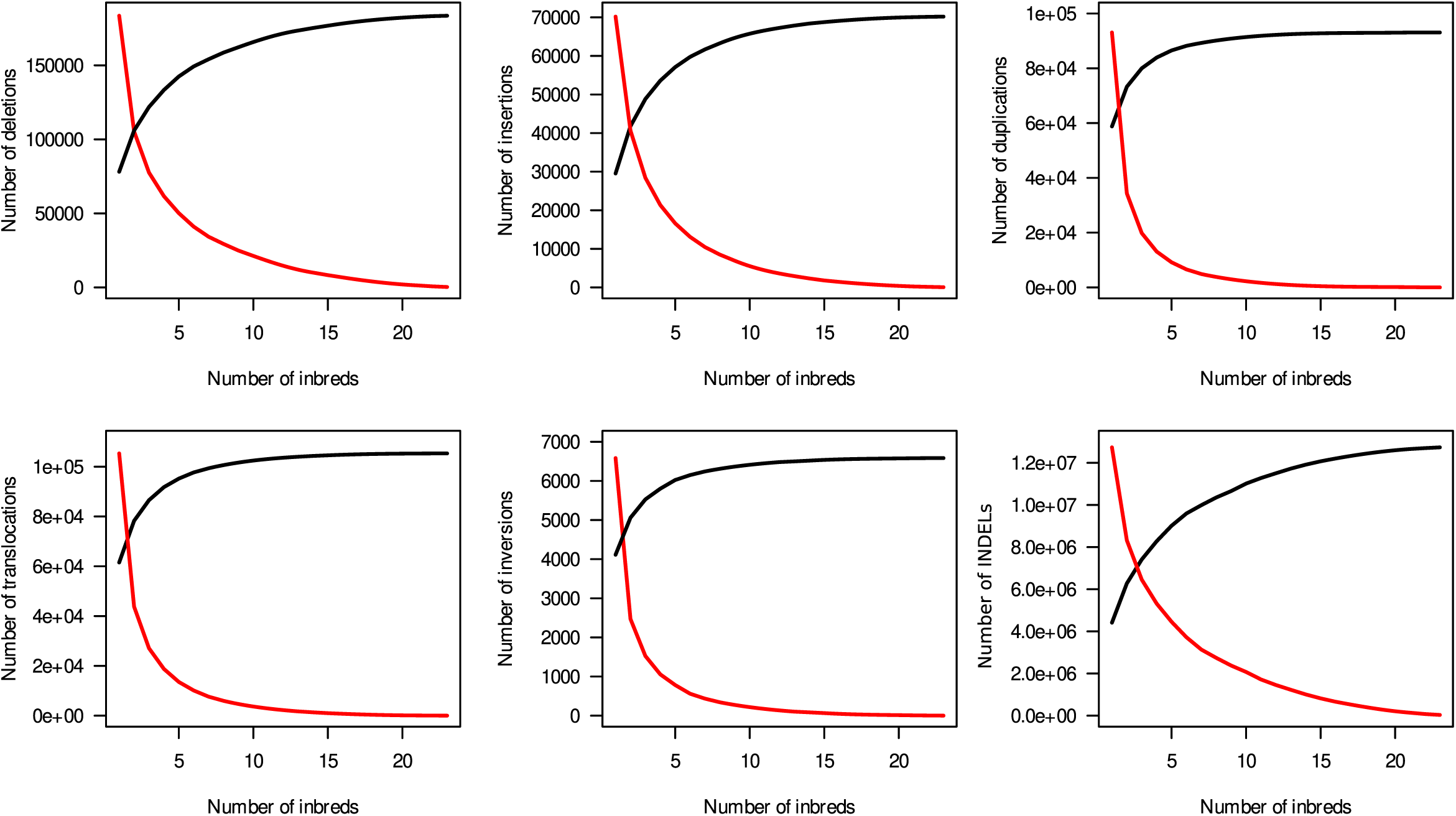
Number of structural variant (SV) clusters for the different types of SV which were detected in at least (red) or no more than (black) the given number of inbreds.

**Fig. S8:**
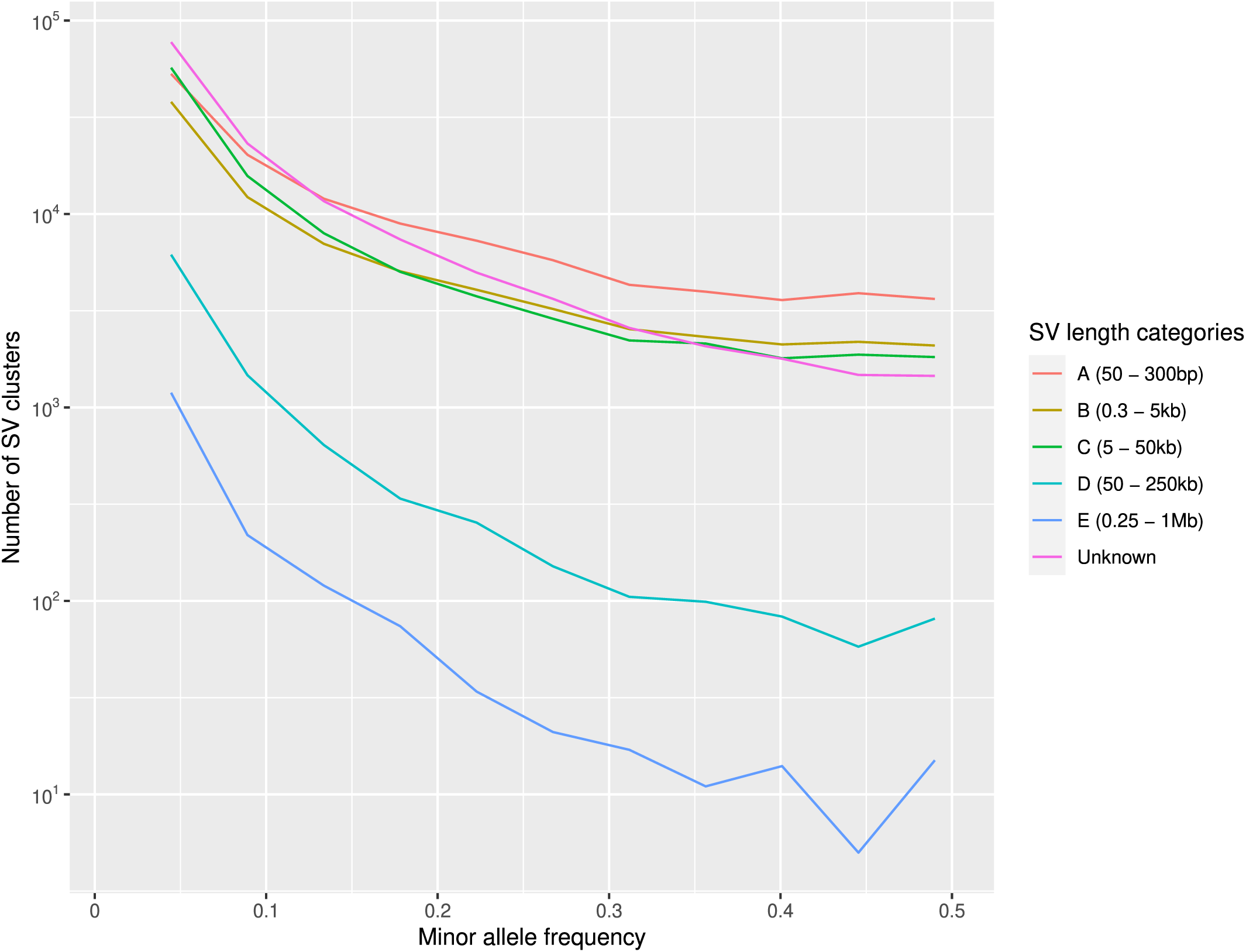
Detection frequencies of structural variant (SV) clusters of different length categories across the 23 barley inbreds.

**Fig. S9:**
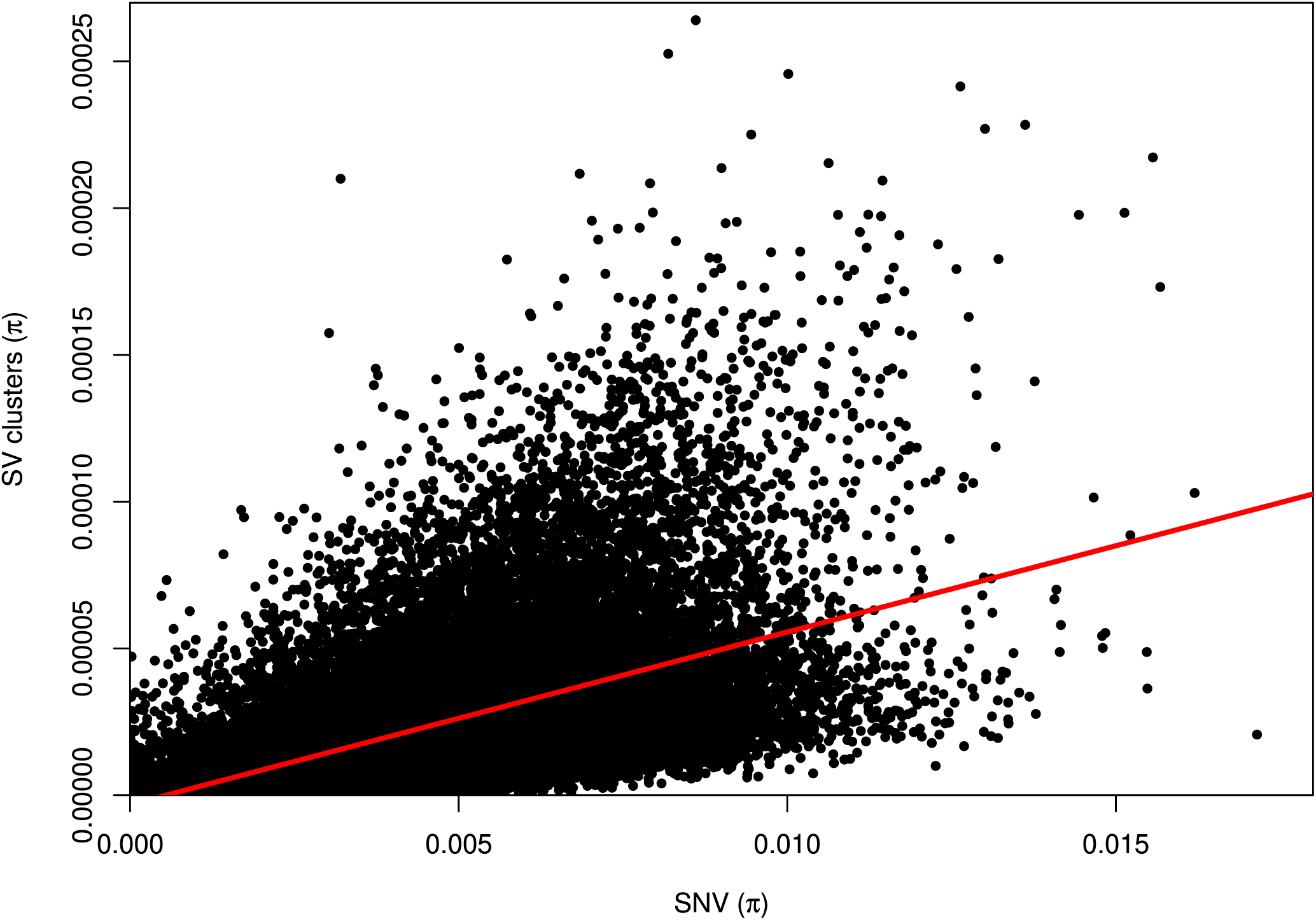
Average genetic diversity (*π*) of SNV and SV clusters across 100kb windows of the genome. The red line indicates the correlation.

**Fig. S10:**
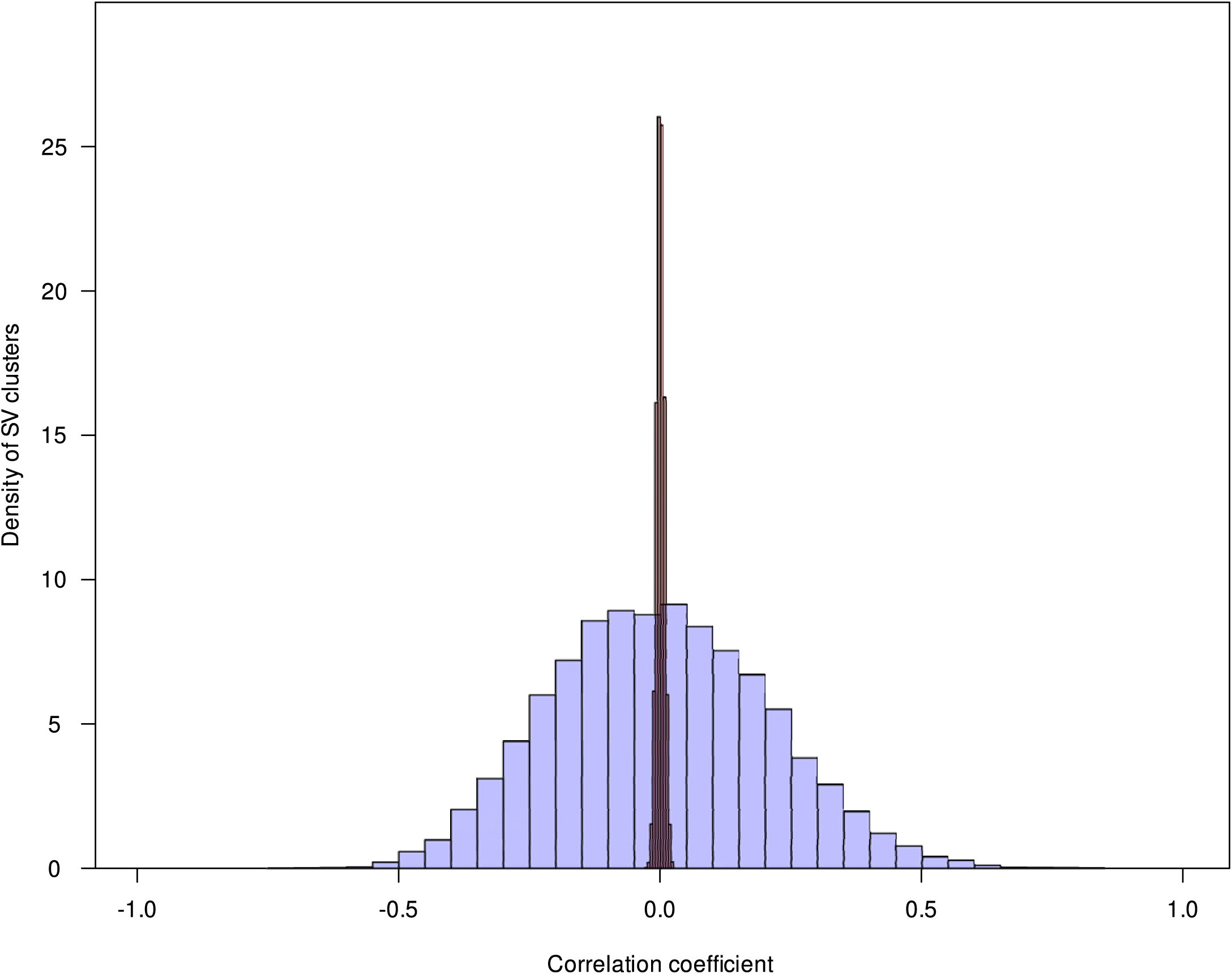
Distribution of correlation coefficients of presence/absence pattern of all structural variant (SV) clusters (deletions, insertions, duplications, inversions) with minor allele frequency *>* 0.15 and the loadings of principal component 1 (19.7%) from a principal component analysis of gene expression data. The blue histogram shows the distribution for the detected SV clusters whereas the red histogram shows the distribution for random SV clusters with identical allele frequency.

**Fig. S11:**
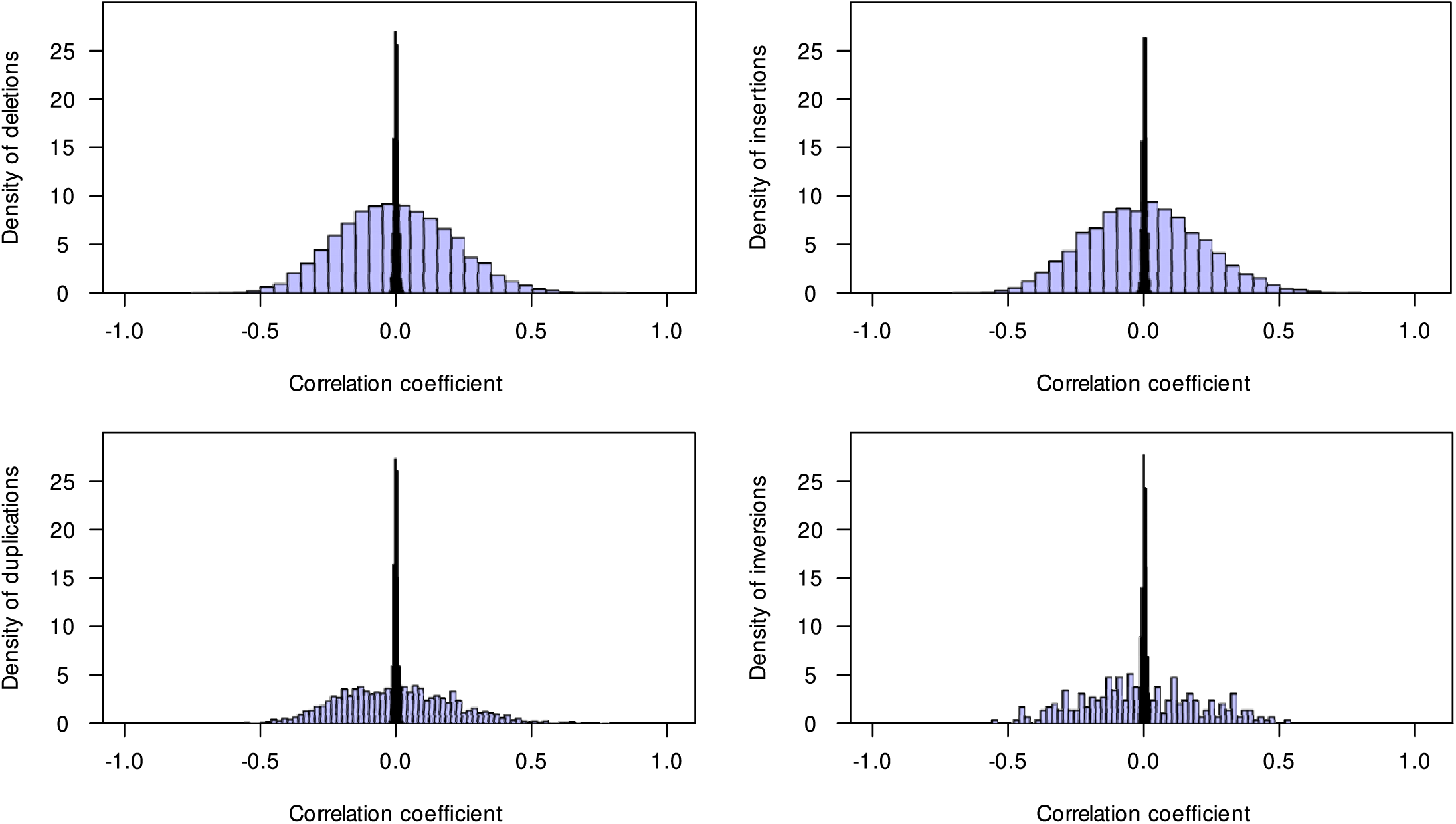
Distribution of correlation coefficients of presence/absence pattern of deletions, insertions, duplications, and inversions with minor allele frequency *>* 0.15 and the loadings of principal component 1 (19.7 %) from a principal component analysis of gene expression data. The blue histogram shows the distribution for the detected SV clusters whereas the red histogram shows the distribution for random SV clusters with identical allele frequency.

**Fig. S12:**
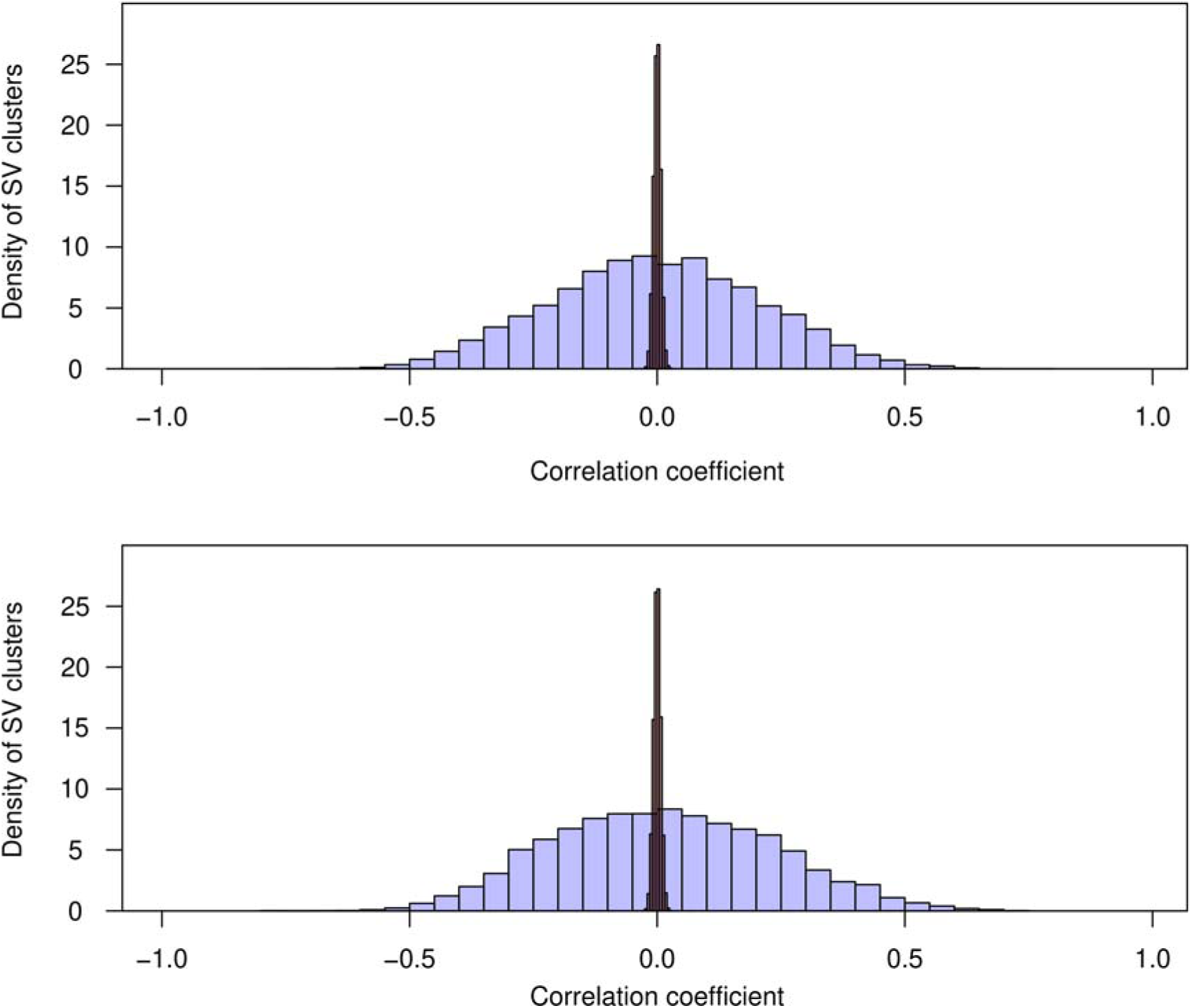
Distribution of correlation coefficients of presence/absence pattern of SV clusters with minor allele frequency *>* 0.15 and the loadings of principal component 2 (8.2 %) (A), and 3 (7.1%) (B) from a principal component analysis of gene expression data. The blue histogram shows the distribution for the detected SV clusters whereas the red histogram shows the distribution for random SV clusters with identical allele frequency.

**Fig. S13:**
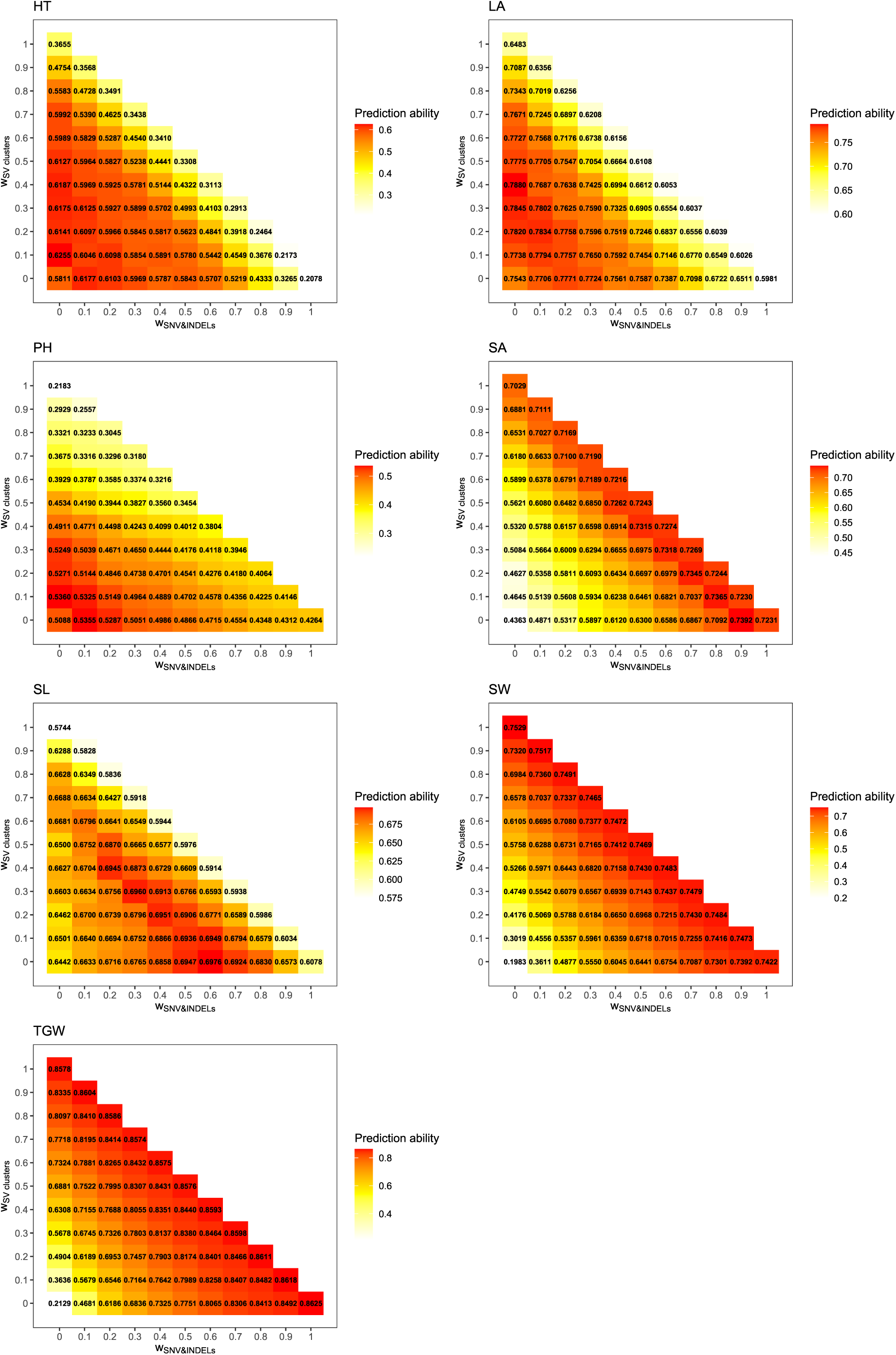
Prediction ability for the seven phenotypic traits heading time (HT), leaf angle (LA), plant height (PH), seed area (SA), seed length (SL), seed width (SW), and thousand grain weight (TGW) from 23 inbreds for 66 combinations of the joined weighted matrices which differ in the weights of three predictors single nucleotide variants (SNV) and small insertions and deletions (2 - 49bp, INDELs, SNV&INDELs, x-axis), structural variant (SV) clusters (y-axis), and gene expression. Plotted values represent medians across 200 cross-validation runs.

